# The mitochondrial calcium uniporter transports Ca^2+^ via a ligand-relay mechanism

**DOI:** 10.1101/2023.06.17.545435

**Authors:** Connie Chan, Chen-Ching Yuan, Jason G. McCoy, Patrick S. Ward, Zenon Grabarek

## Abstract

The mitochondrial calcium uniporter (mtCU) is a multicomponent Ca^2+^-specific channel that imparts mitochondria with the capacity to sense the cytosolic calcium signals. The metazoan mtCU comprises the pore-forming subunit MCU and the essential regulator EMRE, arranged in a tetrameric channel complex, and the Ca^2+^ sensing peripheral proteins MICU1-3. The mechanism of mitochondrial Ca^2+^ uptake by mtCU and its regulation is poorly understood. Our analysis of MCU structure and sequence conservation, combined with molecular dynamics simulations, mutagenesis, and functional studies, led us to conclude that the Ca^2+^ conductance of MCU is driven by a ligand-relay mechanism, which depends on stochastic structural fluctuations in the conserved DxxE sequence. In the tetrameric structure of MCU, the four glutamate side chains of DxxE (the E-ring) chelate Ca^2+^ directly in a high-affinity complex (site 1), which blocks the channel. The four glutamates can also switch to a hydrogen bond-mediated interaction with an incoming hydrated Ca^2+^ transiently sequestered within the D-ring of DxxE (site 2), thus releasing the Ca^2+^ bound at site 1. This process depends critically on the structural flexibility of DxxE imparted by the adjacent invariant Pro residue. Our results suggest that the activity of the uniporter can be regulated through the modulation of local structural dynamics.

A preliminary account of this work was presented at the 67^th^ Annual Meeting of the Biophysical Society in San Diego, CA, February 18-22, 2023

## Significance

The mitochondrial calcium uniporter is a multi-subunit protein complex that integrates cellular Ca^2+^ signaling with mitochondrial energy output. It is a highly selective, high-conductance Ca^2+^ channel. The structure of the uniporter is known, but the mechanism of its regulation is controversial. Our work elucidates the mechanism of the uniporter’s Ca^2+^ conductance and has broad implications for understanding the function of ion-selective channels. The ligand-relay mechanism that we propose offers insights into the persistent in the ion channel field question of how the specificity of ion recognition is reconciled with a nearly diffusion-limited rate of their conductance. Unexpectedly our work reveals the fundamental role of the local structural dynamics in the uniporter’s function and suggests an alternative mechanism of its regulation.

## Introduction

Transient spikes in the concentration of cytosolic Ca^2+^ regulate a variety of cellular functions. A small fraction of each Ca^2+^ spike exceeding a certain activation threshold is taken up by the mitochondria to regulate multiple processes adjusting the organelle’s metabolic state to the cell’s energy requirements. Mitochondria uptake Ca^2+^ *via* the uniporter (mtCU), a highly selective, high-conductance Ca^2+^-activated channel (1) residing within the mitochondrial inner membrane (2–4). The uniporter contains a pore-forming subunit MCU (3, 5), a metazoan-specific single-pass trans-membrane subunit EMRE (6), and the EF-hand Ca^2+^-binding peripheral proteins MICU1-3 (2–4, 7).

The atomic resolution structures of MCU from several species have been reported. The first NMR structure of *C. elegans* MCU defined the main features of the channel but had an incorrect symmetry (8). Subsequently, several groups reported the cryo-EM and X-ray structures of MCU from fungi (9–12), showing that the channel is built of four MCU monomers, each contributing the second of its two transmembrane helices (TM1, TM2) to the Ca^2+^ conducting pore. The only part of MCU that is accessible from the intermembrane space (IMS) is a stretch of ∼14 residues, here referred to as the TM-linker, which comprises the C-terminal turn of TM1, a two-residue connector in an extended conformation, and the N-terminal turn of TM2, (residues 252-265 of human MCU (HsMCU)). Included in the TM-linker is the conserved DxxE sequence, which is indispensable for MCU function (3, 13). The four glutamate sidechains of DxxE – one from each monomer –referred to as the E-ring or E-locus, bind Ca^2+^ tightly and constitute the selectivity filter (14). The role of the four aspartates, the D-ring or D-locus, is less clear. In the structure of HsMCU complexed with EMRE (15), the N-terminal segment of EMRE stabilizes the juxtamembrane loop (JML) of one MCU monomer, whereas its single transmembrane helix interacts with the TM1 of an adjacent MCU monomer. Hence, the four EMRE monomers brace the MCU tetramer, contributing to the stability of the complex. The IMS-facing highly conserved C-terminal poly-Asp tail of EMRE has not been resolved. Recent cryo-EM studies yielded nearly atomic resolution structures of reconstituted protein complexes containing all the uniporter subunits (16–19). They show a dimeric assembly of MCU/EMRE tetramers connected by their N-terminal domains on the matrix side and bridged by two MICU1-MICU2 heterodimers on the IMS side. These structures suggest that the subunit stoichiometry of the uniporter is one MICU1-MICU2 heterodimer for an octamer consisting of four MCU and four EMRE monomers. However, it is unclear if such a complex represents the native state of mtCU, considering there have been no reports of successful *in vitro* reconstitution of the uniporter’s full functionality.

Various lines of experimentation have led to contradicting proposals about the mechanism of uniporter regulation. Early observations indicated that MICU1 regulates the calcium activation threshold, i.e., the lowest concentration of free Ca^2+^ at which a net influx into the matrix can be observed (2, 4, 20, 21). Deletion of MICU1 in a model cell line lowered the threshold significantly. Thus, MICU1 has been dubbed “the gatekeeper.” It also became clear that MICU1 performs its function not alone but in tandem with MICU2 (7). The two proteins form a heterocomplex that binds Ca^2+^ cooperatively with a high affinity consistent with the uniporter’s activation threshold (22). Strong support for the gatekeeping hypothesis came from experiments in which the Ca^2+^ binding sites of MICU1 or MICU2 were disabled, thus locking the complex in the apo-conformation and causing loss of the Ca^2+^ conductance (7, 23). In contrast to these studies, no apparent Ca^2+^ threshold or a gating function was found in the stopped-flow experiments on rat ventricular mitochondria (24). Also inconsistent with the channel-blocking function of apo-MICU1 appeared the Ru360 sensitive Na^+^ conductance observed in the patch-clamp experiments on mitoplasts in Ca^2+^-free conditions (25). This result, combined with the observation that mitoplasts from wild-type cells had higher Ca^2+^ uptake activity than those from MICU1 knockout cells, led the authors to the conclusion that MICU1 does not inhibit the uniporter but increases its activity at high [Ca^2+^] (25). This proposal was contradicted in two recent reports showing that the lack of Na^+^ current inhibition was attributable to the loss of MICU1 during the mitoplast preparation (26, 27). These studies reaffirmed the inhibitory role of MICU1, but left open the possibility that the composition and, effectively, the function of mtCU might be variable depending on the cell or tissue type or that it could be modulated dynamically in response to physiological cues.

It remains unclear what constitutes the putative “gate” that controls the Ca^2+^ conductance of mtCU. In one study, a transmembrane mechanism involving the Ca^2+^ binding to a putative inhibitory sensor in the matrix-located N-terminal domain of MCU was postulated (28). A more direct pore-occlusion mechanism was proposed based on the cryo-EM structures of reconstituted uniporter complexes. In these structures, the channel appears to be physically blocked by the Ca^2+^-free MICU1/MICU2 heterodimer (16–19). Since no such occlusion was found in the presence of Ca^2+^ it was proposed that Ca^2+^ binding induces the MICU1/MICU2 heterodimer to dissociate from the channel’s pore, permitting the conductance. While this model appears attractive, it lacks a satisfactory mechanistic explanation for how Ca^2+^ binding to the MICU1/MICU2 heterodimer reduces its affinity for the MCU/EMRE tetramer. The critical requirement of metazoan MCU for EMRE inspired a proposal that the juxtamembrane loop (JML) of MCU - which makes extensive contacts with the N-terminal segment of EMRE - might work as a gate (15, 29, 30). However, in all currently available MCU structures, this putative luminal gate is either open or unstructured. The MCU pore is open throughout except at the selectivity filter, the E-ring of DxxE, where a tightly bound Ca^2+^ blocks the passage. The dissociation constant for Ca^2+^ at this site is reportedly less than 2 nM (1). The origins of such an exceptionally strong binding, and its consequences for the channel’s conductivity and regulation, require elucidation.

In the present study, we performed an extensive analysis of MCU sequence conservation, structure, and dynamics, focusing on the TM-linker region. We show that TM-linker has an irregular structure that is highly dynamic on a nanosecond timescale and that this conserved feature, imparted primarily by the proline residue located next to the DxxE motif, is essential for the Ca^2+^ uptake activity of MCU. Applying hexaamminecobalt(III) as a stable structural analog of a hydrated Ca^2+^ ion in molecular dynamics simulations revealed the details of the Ca^2+^-ligand interactions in the channel’s pore. We propose that the key step in the Ca^2+^ uptake by MCU is the transient sequestration of a hydrated Ca^2+^ via the aspartate side chains of the D-ring of DxxE. It facilitates the dissociation of the glutamate side chains of the E-ring from their direct, mostly bidentate interaction with the Ca^2+^ ion, permitting its release into the matrix. Hence, we propose that the glutamate side chains of the selectivity filter of MCU play an active role in transferring Ca^2+^ between the two loci of DxxE in a process we call a “ligand-relay mechanism.”

## Results

### Sequence conservation in MCU highlights invariant features essential for the function

We have compiled an extensive library of MCU sequences and analyzed them in terms of conservation. Previously, putative MCU homologs were identified by the presence of the essential DxxE sequence, termed DIME motif, flanked by two transmembrane helices (TM1,TM2) and the coiled-coil segments (13). Here, we relied on the database sequence annotations consistent with these criteria. The set of retrieved sequences was curated manually by removing duplicates and sequences with large deletions or insertions in the region corresponding to the transmembrane domain of HsMCU. Also removed were excessively long sequences (>800aa), suggesting a structurally distant analog of uncertain function or a sequencing error. Sequences annotated as MCUb were also removed. The final library contained 2495 MCU sequences distributed among *metazoa*, *viridiplantae,* and *fungi* (SI Appendix Fig. S1). A confounding issue with the curation of our dataset was the lack of a clear distinction between ancient MCU and its inhibitory paralog MCUb. In humans, MCU and MCUb share ∼50% sequence identity. A functionally distinctive feature is the substitution of two charged residues in the last turn of TM1 (R252 and E257 of HsMCU) with hydrophobic residues (W237 and V242, respectively, in HsMCUb), substitutions known to inhibit calcium uptake (31). It is not known what other sequence features distinguish MCUb from MCU.

Alignment of the 2495 MCU sequences with the program MUSCLE (MUltiple Sequence Comparison by Log-Expectation) (32) reveals extreme variability in the N- and C-terminal segments of MCU, featuring large insertions and deletions (Supplement 2), suggesting little evolutionary pressure for the matrix-located parts of MCU, or unknown functional adaptations. In contrast, the central region of the sequence (residues 170-346 of HsMCU), comprising the membrane-spanning helical domain of MCU previously found sufficient for the Ca^2+^ uptake activity in the presence of EMRE (8), is well conserved. We quantified the sequence conservation in this region using ConSurf (33). Not surprisingly, well conserved is the transmembrane helix 2 (TM2), which comprises the channel’s pore, as well as the entire TM-linker region (Fig. 1). The evolutionary conservation of the DxxE motif in the first turn of TM2 is well understood, given its critical role in MCU function (13). Substitutions of either D261 or E264 with Ala greatly reduce or eliminate, respectively, the Ca^2+^ uptake activity of HsMCU (3). In our MCU sequence library, we identified 14 variants at the D-locus (substitutions with E, N, S, H, G, or A) but only one variant at the E-locus. In the MCU tetramer, the four Asp and Glu sidechains of the DxxE motif form distinctive rings of carboxylates, each capable of binding a divalent metal ion independent of the other (34). The E-ring binds Ca^2+^ tightly and serves as the selectivity filter (14). The four aspartates of the D-ring are at a distance suitable for accommodating a hydrated Ca^2+^ ion (11). The conserved Trp on the N-terminal side of DxxE is thought to stabilize the selectivity filter via the sidechain hydrogen bond interactions with the E-ring (9). Among the 2495 sequences in our library, we found only one variant at this position, a substitution with Arg in *Tuber magnatum* MCU. The functional significance of high sequence conservation in other positions of the TM-linker, especially of Pro265, of which we found only two variants, a substitution with Gln in *Eragrostis curvula* MCU or with Ala in *Stentor coeruleus* MCU, is unclear.

**Figure 1.**
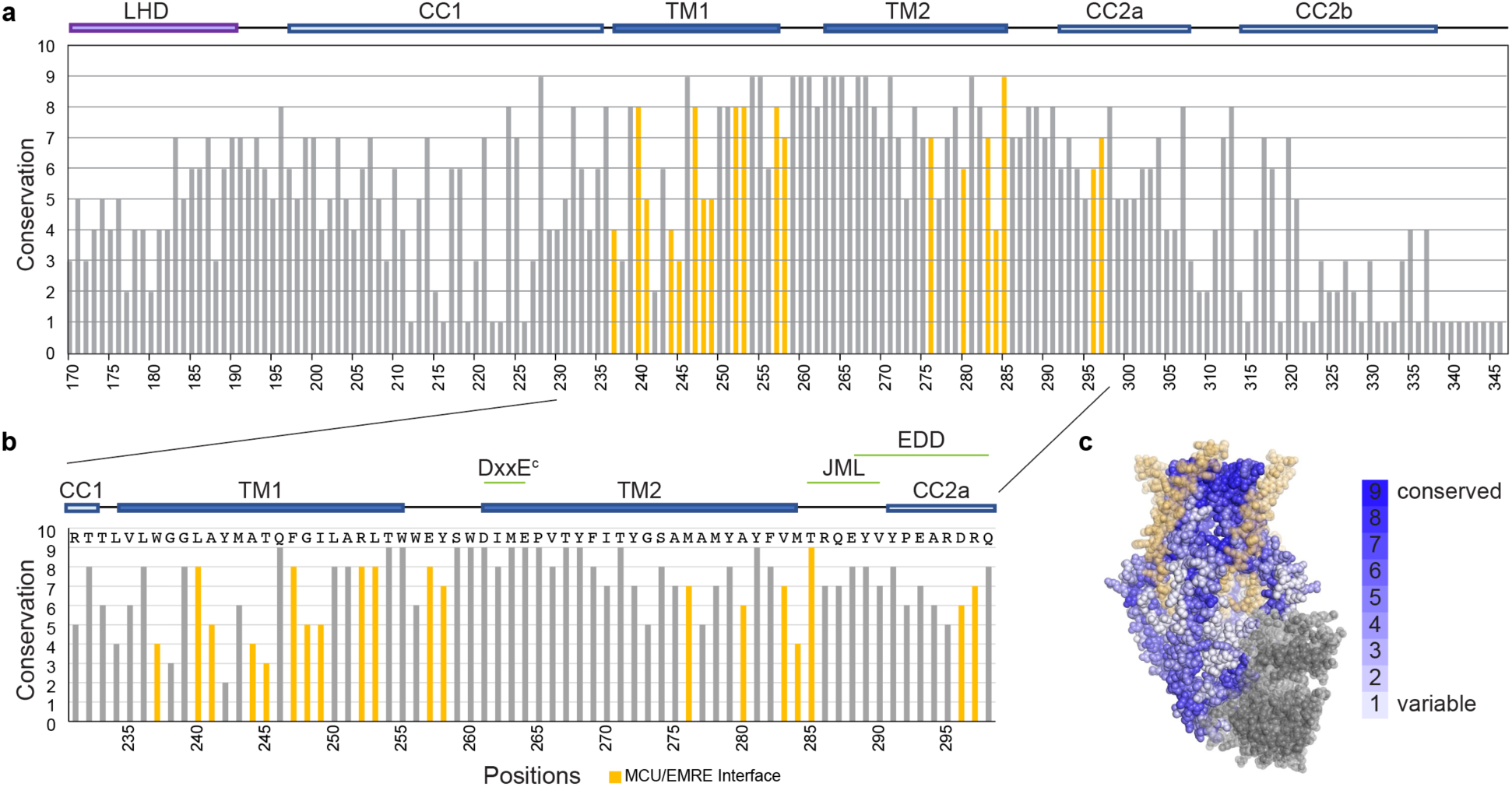
Amino acid sequence conservation in MCU. **A** – The sequence conservation score, as defined by ConSurf, for the helical domain of MCU within a library of 2495 sequences. The highly variable N-terminal region is not shown. The residue numbering corresponds to the human MCU sequence. The abbreviations denote the structural elements: TM1, TM2 – transmembrane helices 1 and 2, respectively, CC1, CC2a, CC2b – the coiled-coil regions, LHD – the linker helix domain, JML – juxtamembrane loop, EDD –EMRE dependence domain. **B** – A zoomed-in view of the transmembrane region of MCU. The sequence of human MCU is shown on top of the graph. The yellow bars identify the residues involved in the interaction with EMRE in the structure of the human MCU-EMRE complex (PDB:6O58) as defined by PDBePISA. **C** – Visualization of the sequence conservation on the structure of the human MCU-EMRE complex. The EMRE subunits are shown in yellow. The N-terminal domain of MCU (shown in gray) was not included in the sequence conservation analysis.

### Flexible secondary structure of the TM-linker region of MCU

Next, we sought to relate the sequence conservation with the secondary structure of MCU. The Ramachandran plot of the transmembrane segment of MCU (residues 234-284 of HsMCU (PDB: 6O58)) shows two distinctly different patterns for the main chain ϕ,ψ dihedral angles. As expected, the ϕ,ψ angles of membrane-embedded sections of TM1 and TM2 are tightly clustered in the region characteristic of α-helical conformation. In contrast, the ϕ,ψ angles of the TM-linker are more scattered (Fig. 2A). For a more intuitive view, we converted the Ramachandran plot into a ϕ^2^,ϕ,ψ^2^ plot (Fig. 2B) and its sequence-ordered representation (Fig. 2C). In the latter plot, the ϕ^2^,ϕ,ψ^2^ values averaged over all eight MCU chains of PDB:6O58 provide a measure of the polypeptide chain’s departure from a canonical α-helical conformation. This plot shows that in addition to the two central residues (Y258, S259), which are in an extended conformation, the last turn of TM1 and the first turn of TM2 depart significantly from the canonical α-helix. We found similar distortions in the TM-linker in other MCU structures that are currently available (SI Appendix Fig. S2).

**Figure 2.**
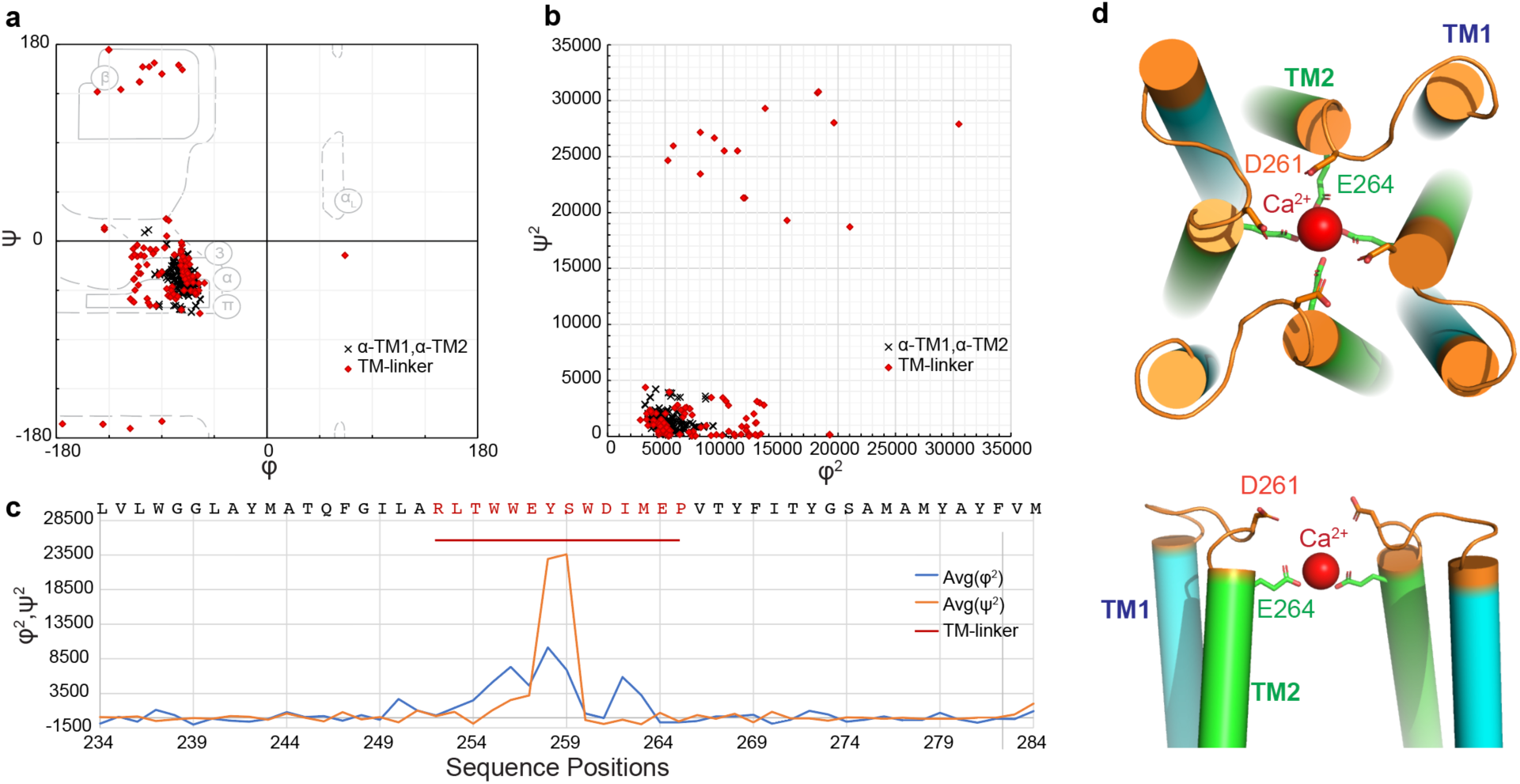
Irregular secondary structure in the TM-linker indicates structural instability. **A** – Ramachandran plot for the transmembrane domain of MCU (residues 234-284). The structure of the human MCU-EMRE complex (PDB: 6O58) was used. The red diamonds indicate the TM-linker region (residues 252-265), and the black symbols show the membrane-embedded parts of TM1 and TM2. Regions of the Ramachandran plot corresponding to various secondary structure types are outlined. **B** – A ϕ^2^,ϕ,ψ^2^ plot highlights the difference in the distribution of main chain dihedral angles between the TM-linker region and the membrane-embedded sections of the TM helices. **C** – Sequence-ordered representation of the ϕ^2^,ϕ,ψ^2^ plot. The average values of ϕ^2^ and ϕ,ψ^2^ for the membrane-embedded section of the transmembrane helices (black symbols in panels A and B) have been subtracted, respectively. **D** – The structure of the TM domain of MCU in two orientations, top view, and side view cross-section. The TM-linker is shown in orange.

Of particular interest is the conformation of the first turn of TM2, which comprises the DxxE motif. The ϕ^2^,ϕ,ψ^2^ values are at baseline for the first two residues (W260 and D261), consistent with α-helical conformation. In contrast, the next two residues (I262, M263) have elevated ^2^values indicating a kink in the helix. The plot returns to baseline for E264 and the remaining part of TM2. The cause of this distortion appears to be the adjacent Pro265, which cannot serve as a hydrogen bond donor for the carbonyl oxygen of Asp261, reducing helix stability. The fact that the Pro residue at this position is virtually 100% conserved (Fig. 1) suggests that the helix distortion in DxxE is essential for MCU function. Due to the kink in the helix, the sidechains of D261 and E264 are more closely aligned with each other than they would be in a canonical α-helix. A different type of helix distortion is present in the last turn of TM1, where the polypeptide chain makes a wider turn akin to a π-helix with the *i,i+5* main chain H-bond pattern. The π-helix is less stable than the canonical α-helix (35). Interestingly, the decrease in the main chain conformational stability may be compensated to some extent by the sidechain hydrogen bond between R252 and E257, which is facilitated in the π-helix but not possible in the α-helix. This structural analysis reveals that the most evolutionarily conserved segment of MCU has an irregular structure, which appears to be flexible. The mechanistic role of the TM-linker flexibility for the Ca^2+^ uptake function of MCU requires elucidation.

### Stochastic dynamics of the TM-linker on a nanosecond timescale

To gain insights into the TM-linker’s structural irregularity, we used molecular dynamics (MD) simulations. The HsMCU-EMRE complex (PDB: 6O58) consisting of four MCU subunits (residues 170-346) and the interacting EMRE subunits (residues 48-96) were placed in a lipid bilayer (Fig. 3A) and a periodic water box supplemented with 150 mM KCl and pH set to 7.0. We also included the single Ca^2+^ ion tightly bound in the E-ring of DxxE (site 1). Following an initial structure equilibration, we ran an unsupervised all-atom MD simulation at 310 K for 10 ns. The atomic coordinates of the assembly were recorded every 0.1 ns for a total trajectory of 100 frames. We found no large-scale structural instabilities during the run, except for a partial dissociation of the N-terminal segment of EMRE from its interaction site on MCU. In the subsequent MD runs, we omitted the EMRE subunits and used only the transmembrane part of MCU (residues 216-310), which was stable during the simulation.

**Figure 3.**
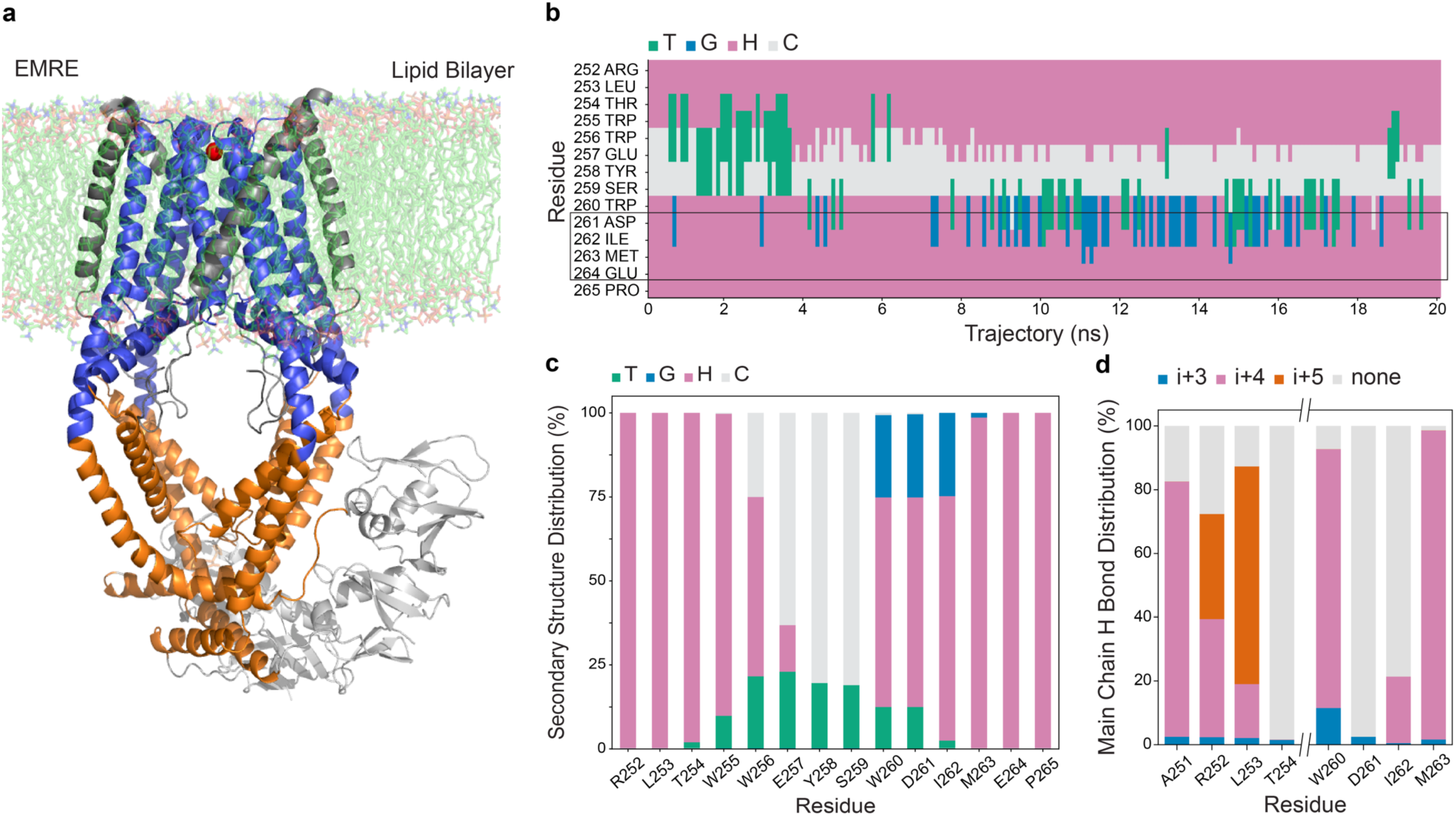
Molecular dynamics simulations confirm the structural instability of the TM-linker region of MCU. **A** – Structure of the human MCU-EMRE complex (PDB:6O58) embedded in a lipid bilayer. The segment used for MD simulations is shown in blue. **B** – Secondary structure fluctuations in the TM-linker region. Color coding for the secondary structure types: Green - turn (T), Blue - 3_10_- helix (G), Pink - α-helix (H), Grey - coil (none of the above) (C). **C** – Secondary structure distribution. **D** – Distribution of the main chain hydrogen bond types in the last turn of TM1 and the first turn of TM2. Color coding for hydrogen bond types: blue - i,i+3 (3_10_-helix), pink - i,i+4 (α-helix), orange - i,i+5 (π-helix), gray - none.

A 20 ns MD simulation of WT-HsMCU indicates that the membrane-embedded parts of TM1 and TM2 retain their α-helical conformation. In contrast, the TM-linker undergoes secondary structure fluctuations on a nanosecond time scale (Fig. 3B, SI Appendix Fig. S3). Analysis of the main chain conformation reveals a mixture of secondary structure types, including α-helix, turn, coil, 3_10_ -helix, and ρε-helix (Fig. 3C,D). There is a significant contribution of the *i,i+5* hydrogen bond pattern characteristic of the ρε-helix conformation for R252 and L253 (Fig. 3D). The entire TM-linker remains highly flexible, featuring frequent switching between various secondary structure types. Importantly, the conformation of the E-ring of DxxE remains stable during the run, and the sidechains of E264 remain engaged in the interaction with Ca^2+^. In contrast, the D-ring (D261) is highly dynamic with respect to both the backbone conformation and sidechain orientation.

As pointed out above, the cause of instability within the DxxE sequence appears to be the adjacent invariant Pro265, whose pyrrolidine ring precludes the formation of a hydrogen bond with the main chain carbonyl oxygen of D261. To test the significance of the missing hydrogen bond on the TM-linker’s structure, we substituted Pro265 with alanine. The MD simulation of the P265A-HsMCU mutant shows a significant increase in α-helix contribution for residues 260-262 and the loss of 3_10_ helix conformation (Fig. 4A,B, SI Appendix Fig. S4)). Interestingly, the P265A mutation also causes a decrease in ρε-helix contribution in the last turn of TM1, suggesting a conformational coupling between the two sections of the TM-linker (Fig. 4C). We analyzed the irregularity of the TM2 helix in more detail. As shown in Fig. 4D, the first turn of TM2 in WT-HsMCU is tilted away from the helix axis, resulting in a distinctive kink. The distortion is limited to D261 and I262, as reflected by a significant elongation of the C_α_-C_α_ distances between these residues and those in *i+3* and *i+4* positions, compared to canonical α-helix (Fig. 4E, SI Appendix Table 1). In the P265A mutant, the C_α_-C_α_ distances are more consistent with α-helical conformation. With the kink significantly reduced, D261 and I263 form the helix-stabilizing hydrogen bonds with A265 and V266, respectively.

**Figure 4.**
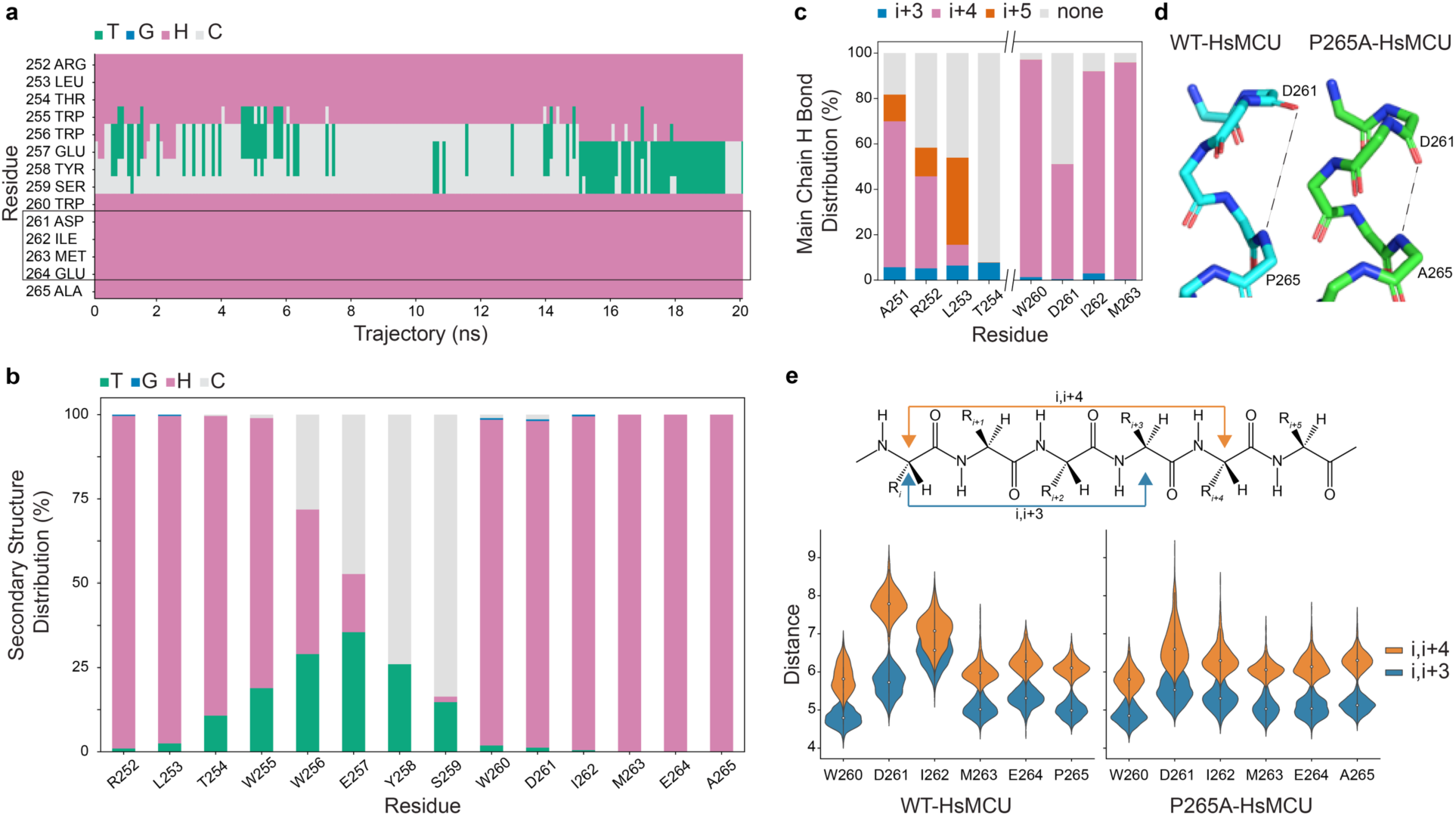
Substitution of P265 with Ala increases the stability of a-helical conformation in the DxxE motif. **A** – Secondary structure fluctuations in the TM-linker. See Fig. 3 for an explanation of the color coding. **B** – Secondary structure distribution. **C** – Distribution of the main chain hydrogen bond types. Refer to Fig. 3 for an explanation of the color coding. **D** – The first turn of TM2 (the DxxE region) in WT-HsMCU has a distinctive kink. The P265A mutation restores the α-helical conformation. **E** – The structural distortion in the DxxE region during MD simulation as measured by the C_α_ distances in i,i+3 and i,i+4 positions. The irregularity of the helical conformation is greatly reduced in the P265A MCU mutant.

**Table 1.** Inhibition of the mitochondrial Ca^2+^ uptake with hexaamminecobalt(III) in permeabilized HEK293T cells expressing various forms of the uniporter. MEAN ± SEM values are shown for n number of measurements.

|  | DKO<br>+WT-HsMCU<br>(n=3) | MCU-KO<br>+WT-HsMCU<br>(n=4) | MCU-KO<br>+D261A-HsMCU<br>(n=3) | MICU1-KO<br>(n=3) |
| --- | --- | --- | --- | --- |
| Uptake rate (A.U.) | 38.7 ± 2.4 | 37.4 ± 5.8 | 1.60 ± 0.01 | 13.6 ± 0.50 |
| IC<sub>50</sub> ($\mu\text{M}$) | 26.7 ± 0.9 | 14.9 ± 0.26 | 210 ± 15.0 | 4.63 ± 0.23 |

### The structural flexibility of the DxxE motif is required for the Ca^2+^ uptake activity of MCU

If the dynamics of the TM-linker is essential for the Ca^2+^ conductance of MCU, then mutations that stabilize this region should be deleterious to the Ca^2+^ uptake activity. The P265A mutation has such a structure-stabilizing effect for the DxxE part of the TM-linker. Hence, we tested the Ca^2+^ conductance of the P265A mutant of HsMCU. We overexpressed this mutant in double-knockout (DKO) HEK293T cells in which the MCU and MCUb genes have been deleted using CRISPR-Cas9 technology (36). We generated two cell lines: DKO-WT-HsMCU and DKO-P265A-HsMCU. We recorded the time course of mitochondrial Ca^2+^ uptake in these cells after permeabilization with digitonin. The WT-HsMCU-expressing cells show robust mitochondrial Ca^2+^ uptake. In contrast, there is no Ca^2+^ uptake in cells expressing the P265A-HsMCU or in the DKO cells (Fig. 5). The P265A mutation completely abolished the Ca^2+^ conductance of MCU. This result, combined with our MD simulations and structural analysis, strongly supports the notion that the structural flexibility of the DxxE motif imparted by the adjacent Pro is essential for the Ca^2+^ conductance of MCU.

**Figure 5.**
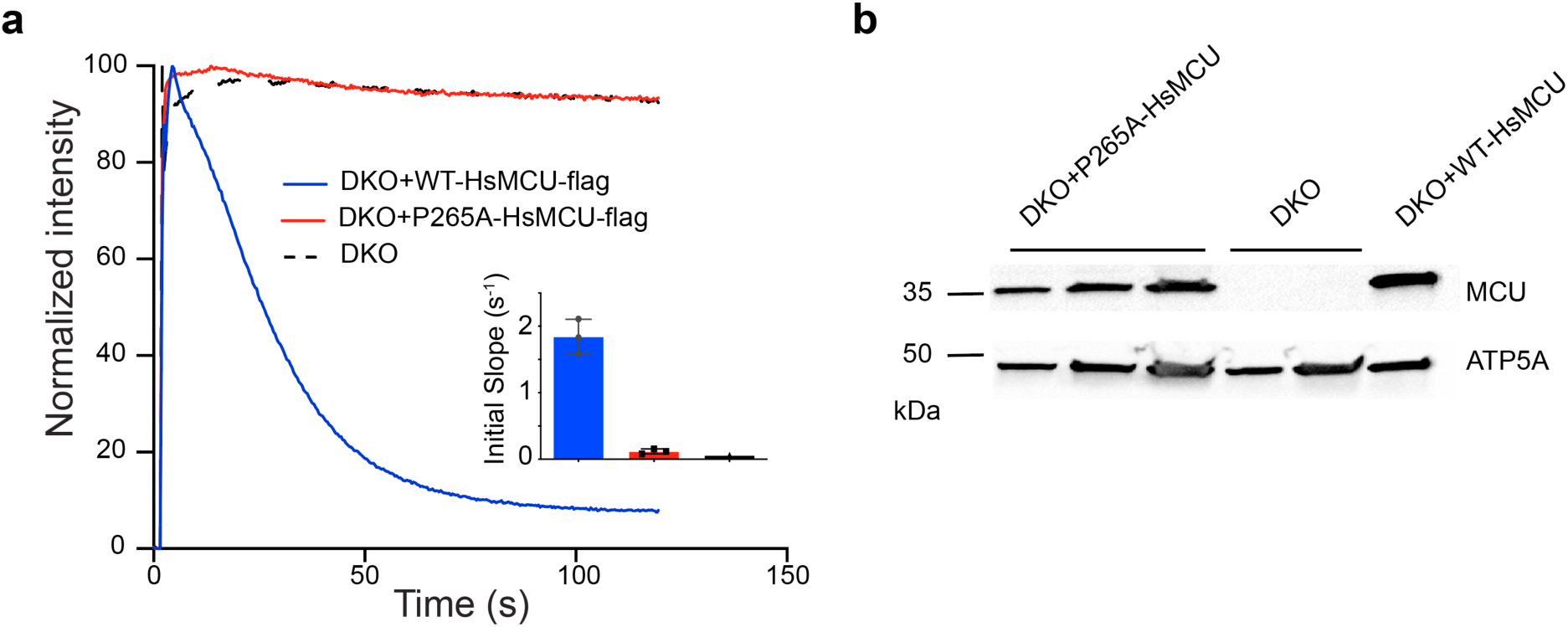
Substitution of the invariant P265 with Ala renders MCU inactive. **A** – Uptake of Ca^2+^ monitored by fluorescence intensity of Calcium Green 5N in permeabilized HEK293T cells expressing the WT-HsMCU or the P265A-HsMCU mutant on the background of a double knockout (DKO) HEK293T cell line in which both MCU and MCUb have been disabled. **B** – Western blot of the corresponding cell lines.

### The critical role of the D-ring in sequestering the hydrated Ca^2+^ ion

Although the requirement for structural flexibility of DxxE is evident, its specific role in the Ca^2+^ conductance of MCU is unclear. The sidechains of D261 are too far apart in the HsMCU tetramer to bind simultaneously to a dehydrated Ca^2+^, and they appear to be somewhat dispensable. Their substitution with Asn in fungal MCU (*Neosartorya fischeri*) has little effect on the Ca^2+^ uptake, and the substitution with Glu improves the uptake (9). In another study, substituting Asp with Ala in *Metharhizium acridum* MCU had a mild effect on the Ca^2+^ conductance (10). It has been proposed that D261 might contribute to the uniporter’s function by interacting with a hydrated Ca^2+^ ion (10, 11). To test this potential role of D261, we utilized hexaamminecobalt(III), ([Co(NH_3_)_6_]^3+^), here abbreviated as NCO, as a stable structural analog of a hydrated Ca^2+^ ion. Although the ionic radius of Co^3+^ is ∼25% smaller than that of Ca^2+^ (0.75 Å and 1.0 Å, respectively (37)), both ligands (NH_3_ and H_2_O) are capable of hydrogen bond interaction with neighboring carboxyls. Furthermore, the octahedral geometry of NCO closely approximates that of a hydrated Ca^2+^ ion ([Ca(H_2_O)_6-8_]^2+^) (38). The advantage of NCO for our studies is the stability of the [Co(NH_3_)_6_]^3+^ complex, in contrast to [Ca(H_2_O)_6-8_]^2+^ in which H_2_O coordinated with Ca^2+^ exchanges rapidly with bulk water. Hence, we set up the MD simulations of MCU and its D261A mutant with a single molecule of [Co(NH_3_)_6_]^3+^ placed within the D-ring of MCU (site 2) in addition to the Ca^2+^ bound in the E-ring (site 1). A 20 ns MD simulation confirmed that NCO is retained within the D-ring due to the formation of numerous H-bond interactions with the sidechain carboxyls of Asp261. In contrast, the D261A-HsMCU mutant could not retain NCO, allowing it to dissociate during the initial 100 picoseconds of the simulation. Clearly, Asp261 of HsMCU is important for the binding of [Co(NH_3_)_6_]^3+^. By inference, these experiments support the view that the D-ring of DxxE serves as an initial binding site for the incoming hydrated Ca^2+^ ion and such transient sequestration, presumably followed by dehydration of [Ca(H_2_O)_6-8_]^2+^ and its transfer to site 1, facilitates the Ca^2+^ conductance.

### Inhibition of the Ca^2+^ uptake of WT and mutant MCU by hexaamminecobalt(III)

To further test the role of D261, we measured the Ca^2+^ uptake rate of the D261A mutant of HsMCU. We have expressed this mutant in HEK293T cells where the MCU gene has been knocked out. We found that the mutation (MCU-KO+D261A-HsMCU) reduces the Ca^2+^ uptake rate to ∼4% of that for the MCU-KO+WT-HsMCU. However, unlike the P265A mutation, the D261A mutation does not abolish the uptake completely (Fig. 6A). Clearly, the aspartate at this position greatly facilitates the Ca^2+^ conductance. Still, it is effective only when the DxxE motif retains its conformational flexibility imparted by the adjacent Pro. It cannot support the Ca^2+^ conductance when DxxE folds into a stable α-helical conformation, as is the case for the DKO+P265A-HsMCU mutant.

**Figure 6.**
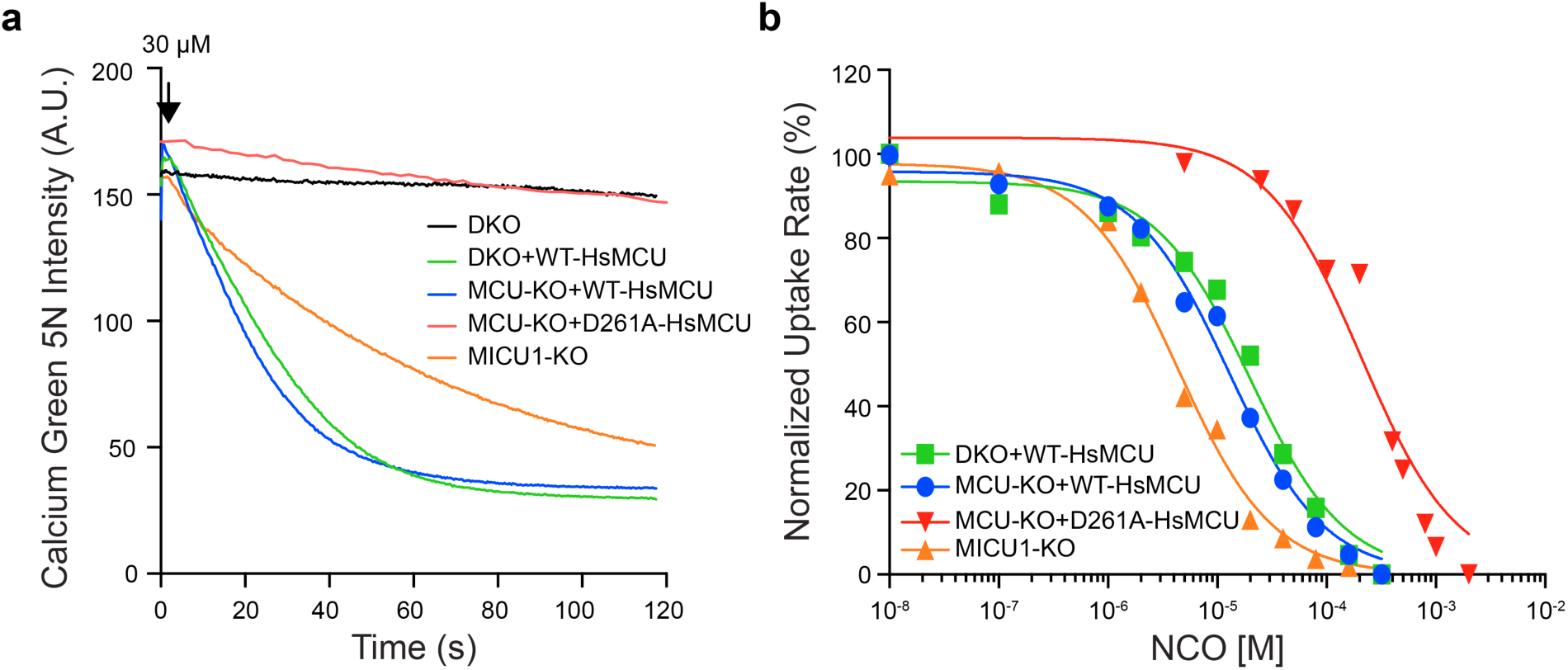
Inhibition of the mitochondrial Ca^2+^ uptake with hexaamminecobalt (III) in permeabilized HEK293T cells expressing various forms of the uniporter. **A** – Representative Ca^2+^ uptake traces in the absence of the inhibitor. **B** – Normalized Ca^2+^ uptake rates as a function of increasing inhibitor concentrations. The lines represent the best fit to a binding isotherm. See Table 1 for the relative Ca^2+^ uptake rates and IC_50_ values.

Our MD simulations provide a vivid illustration of the mechanism by which NCO can inhibit MCU. We verified this by measuring the rates of Ca^2+^ uptake as a function of increasing concentrations of NCO in permeabilized HEK293T cells expressing WT-HsMCU or its variants. We find that micromolar concentrations of NCO inhibit the Ca^2+^ uptake in HEK293T cells (Fig. 6, Table 1), consistent with the early report of Ca^2+^ uptake inhibition by NCO in rat liver mitochondria (39) and a recent study in HeLa cells (40). Interestingly, the very slow Ca^2+^ uptake by the MCU-KO+D261A-HsMCU mutant can also be inhibited by NCO, but at more than tenfold higher concentrations (Table 1). The IC_50_ for cells expressing MCU/EMRE alone (MICU1-KO cells) is roughly fivefold lower than that for the WT-HsMCU expressing cells. This effect is consistent with the observation that Ru360, a commonly used MCU inhibitor, is also more potent in the absence of MICU1 (41).

Altogether, these results support the view that the D-ring’s role in the Ca^2+^ conductance of MCU is to sequester the hydrated Ca^2+^ transiently. They also validate NCO as a stable analog of a hydrated Ca^2+^ ion.

### The mechanism of Ca^2+^ release from the selectivity filter of MCU

Having verified that NCO engages the aspartates of the D-ring of DxxE, we analyzed the effect of this compound on the E-ring’s interaction with Ca^2+^ in the WT-HsMCU and the P265A mutant. The MD simulations showed that both proteins similarly bind NCO, primarily through the hydrogen bonds between the carboxyl side chains of D261 and the NH_3_ groups of NCO. However, in WT-HsMCU, NCO tends to move slightly closer to the bound Ca^2+^ having a profound effect on the E-ring’s interaction with Ca^2+^. In Fig. 7, the status of the E-ring for each of the 200 frames of a 20 ns MD simulation is represented by the sum of distances between the bound Ca^2+^ and the eight E264 side chain oxygens (ΣCa-O). For the WT-HsMCU, there is a much greater range of Ca-O distances, with a dramatic increase in their values at short Ca^2+^-Co^3+^ distances compared to P265A-HsMCU (Fig. 7A,B). The significance of this observation becomes apparent when one considers that the optimal Ca^2+^-oxygen bond length is ∼2.4 Å. When all four glutamates of the E-ring are engaged in a bidentate interaction with Ca^2+^, resulting in a coordination number (CN) of 8, the ΣCa-O parameter has a minimum value of ∼19.2 Å. An increase in ΣCa-O signifies the weakening of the binding due to a progressive breaking of the Ca^2+^-oxygen bonds. The data in Fig. 7 indicate that WT-HsMCU most frequently has CN = 7, and during the simulation, there is a large population of complexes with as few as 4 or 3 Ca^2+^-oxygen bonds remaining. Clearly, NCO causes a dramatic weakening of the Ca^2+^ interaction with the E-ring of MCU. This effect is absent in the P265A-HsMCU mutant (Fig. 7B), for which CN 6 and frequently CN = 8, suggesting increased stability of the Ca^2+^-E-ring interaction. The loss of Ca^2+^ conductance upon the P265A mutation at an equivalent position in *Neosartorya fischeri* MCU has been previously reported and tentatively attributed to reduced stability of the Ca^2+^ binding site (9). Our results do not corroborate this prediction.

**Figure 7.**
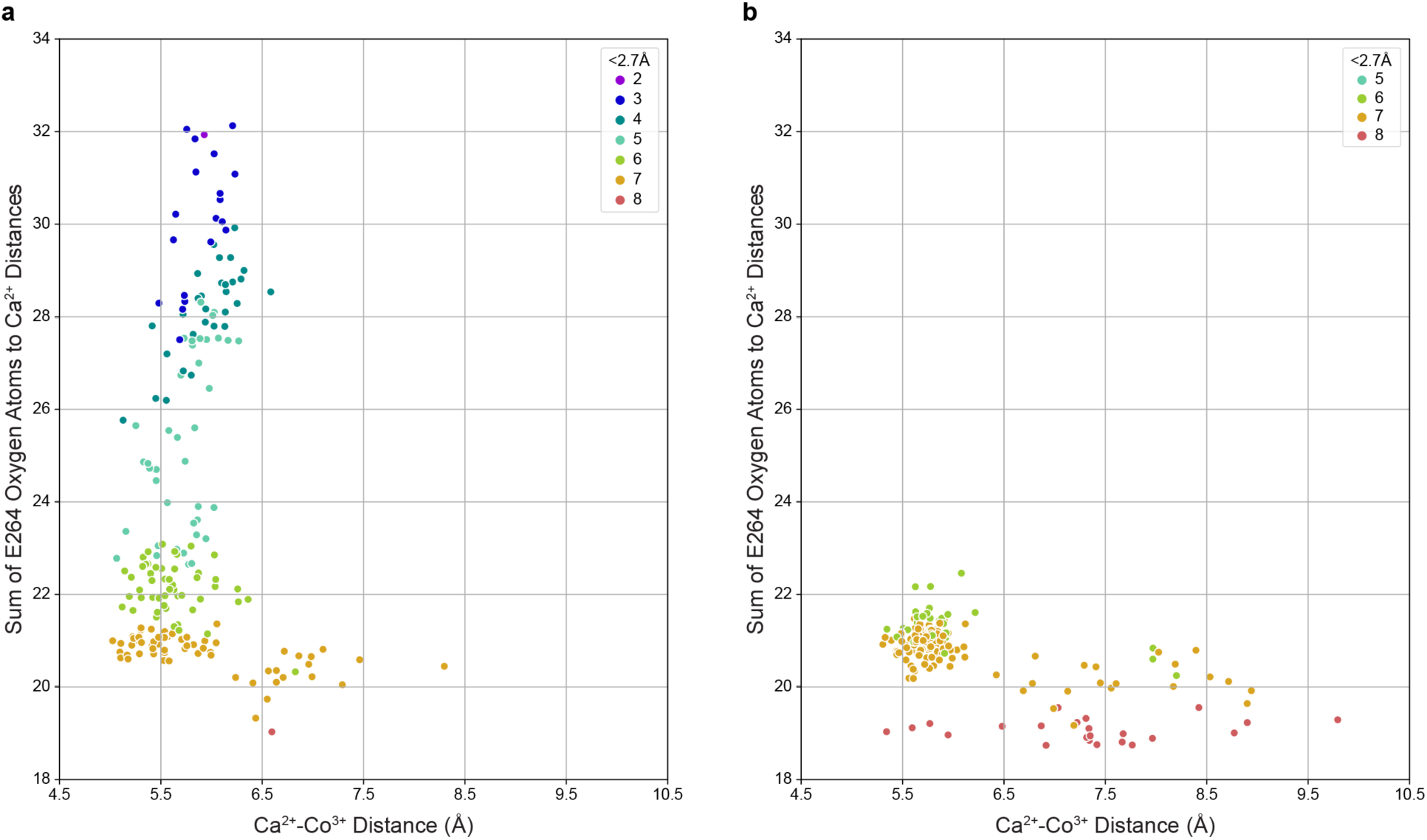
The effect of NCO on the E-ring’s interaction with Ca^2+^ in WT-HsMCU and P265A-HsMCU. Each of the 200 data points represents one frame of the 20 ns MD simulation. The ordinate axis shows the sum of distances between the side chain oxygen atoms of E264 and the bound Ca^2+^. The abscissa is the distance between the Co^3+^ ion of NCO and the Ca^2+^ ion. Increasing values on the y-axis indicate a loss of Ca^2+^-oxygen interaction, weakening the binding. The color coding of the data points (rainbow colors from red to purple) indicates the number of oxygen atoms interacting with the Ca^2+^ (the cutoff distance for the Ca^2+^-oxygen bond is 2.7Å). Note the dramatic weakening of the E264-Ca^2+^ interaction with the approaching NCO for the WT-HsMCU (panel **A**) and the absence of this effect for the P265A mutant (panel **B**).

While the cumulative distance parameter ΣCa-O provides a simple measure of the state of the E-ring, a structural interpretation of this result is needed. We analyzed the distances between the bound Ca^2+^ ion and the two carboxylate oxygen atoms of each E264. The histogram of these values for the WT-HsMCU features five distinctive peaks (Fig. 8). The most prominent peak centered at ∼4.8 Å represents the bidentate COO^-^-Ca^2+^ coordination (the type A interaction in Fig. 8 and Suppl. Fig. 5). The second most abundant peak centered at ∼6.7 Å is the monodentate COO^-^-Ca^2+^ interaction (type B in Fig. 8). Frequently, the monodentate Ca^2+^ interaction permits the side chain E264 to engage in a bridging H-bond interaction with NCO (the subset represented by the gray bars in Fig. 8). Interestingly, for the P265A mutant the distribution is shifted toward type A interaction, i.e., the P265A mutation stabilizes the bidentate Ca^2+^ coordination. The third broad peak for the WT-HsMCU, the type C interaction (∼10.5-12.0 Å), represents the state in which E264 is detached from Ca^2+^, and both its side chain COO^-^ oxygens are H-bonded to NCO. We found no examples of this type of interaction in our MD simulations of P265A-HsMCU. These results explain the loss of activity in the P265A-HsMCU mutant. The reduced main chain flexibility of DxxE precludes the side chains of E264 from fully engaging with the incoming hydrated Ca^2+^ ion sequestered in the D-ring of DxxE. The P265A mutation breaks the communication between the E-ring and the D-ring of DxxE.

**Figure 8.**
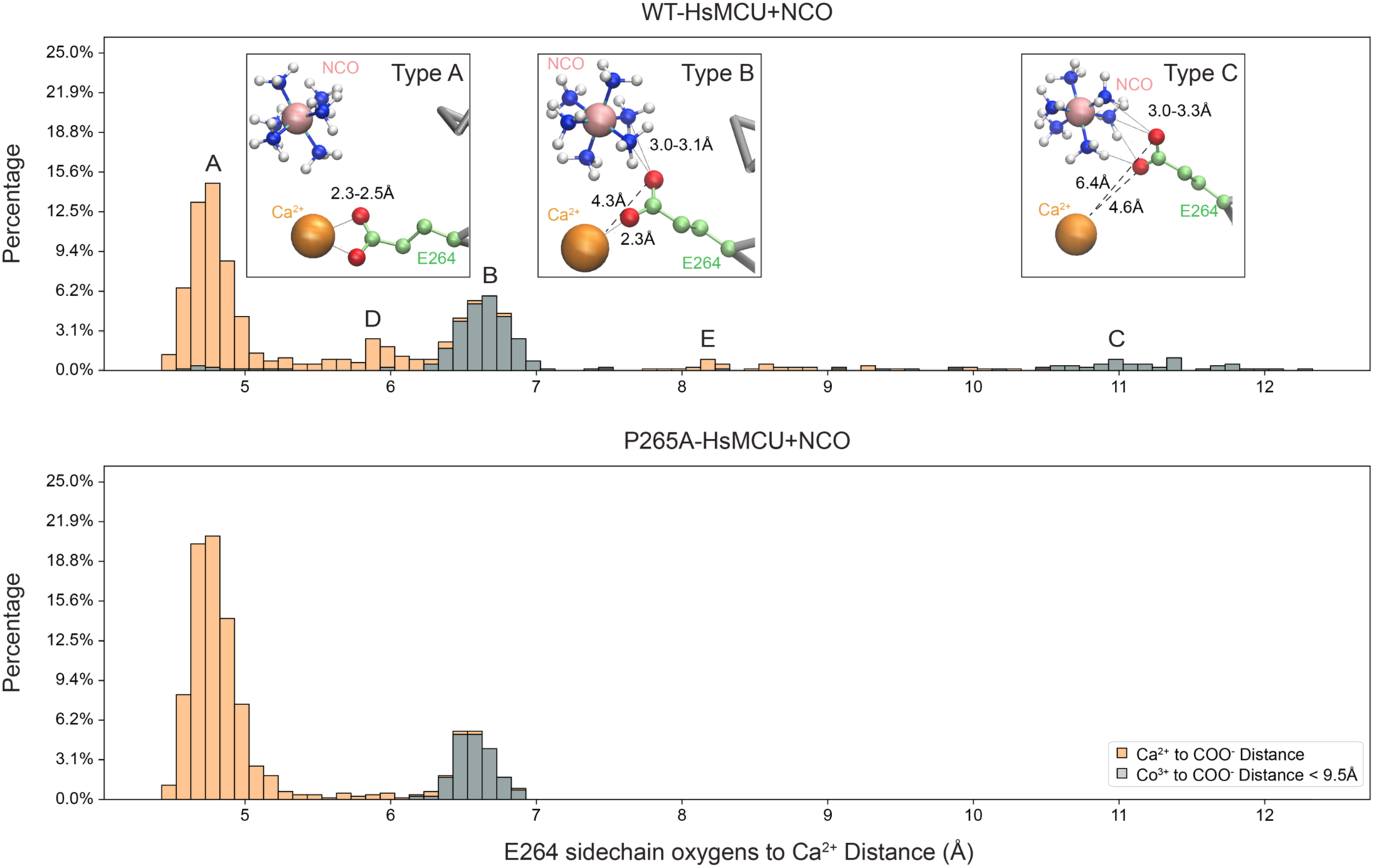
The flexibility of the ligand-calcium interactions in the E-ring of MCU. The relation of the Glu264 side chains to the bound Ca^2+^ is characterized by the sum of the distances of its carboxyl oxygen atoms to Ca^2+^. Five types of interactions are identified: **Type A** – bidentate Ca^2+^ chelation. Both COO^-^ oxygen atoms are bound to the Ca^2+^ ion. **Type B** – monodentate/bridging interaction. One COO^-^ oxygen interacts with Ca^2+,^ while the other connects to NCO via a hydrogen bond. **Type C** – detached/H-bonded to site 2. E264 side chain is detached from Ca^2+^ and interacts entirely with NCO. **Type D** – monodentate/free. One oxygen is engaged in monodentate interaction with Ca^2+^, while the other is free. **Type E** – detached. Neither of the oxygens is involved in binding Ca^2+^ or NCO. The three most populated types are illustrated in the insets. See SI Appendix Fig. S5 for more representative snapshots of the various interactions. Gray bars in the histogram represent the subset of the E264 side chains for which at least one oxygen atom is positioned within <9.5 Å from the Co^3+^ ion, i.e., sufficiently close to engage in a hydrogen bond interaction with an NH_3_ group of NCO. Note the absence of Type C interactions in the P265A-HsMCU mutant.

## Discussion

The important finding of this work is that the calcium conductance of mtCU depends critically on the structural dynamics of the DxxE motif of MCU. This property, imparted primarily by the adjacent invariant Pro residue, appears to be required for the ligand-facilitated communication between two Ca^2+^ ions within the pore of the channel, one trapped in the selectivity filter (the E-ring) and one transiently sequestered in a hydrated form in the D-ring. Our molecular dynamics simulations employing a stable analog of the hydrated Ca^2+^ ion reveal the details of this process.

In a key 2004 publication, the Clapham group reported the electrophysiological characteristics of the uniporter. They documented the exceptional Ca^2+^ specificity of this channel and its potential for an enormous ion-carrying capacity (1). The estimated unitary ion flux at 105 mM CaCl_2_, and V_m_= -160 mV is 5 x 10^6^ Ca^2+^ s^-1^ corresponding to the ion’s dwell time in the pore of less than 2×10^-7^s, a nearly diffusion-limited transport rate. Paradoxically, the same work revealed an extremely high-affinity Ca^2+^-binding site (K_d_ ≤ 2 nM, estimated from the Ca^2+^ block of Na^+^ conductance at 150 mM NaCl). This high-affinity site is now recognized to reside in the selectivity filter of the channel (14). Considering the 2 nM equilibrium dissociation constant and with a conservative estimate for the Ca^2+^ binding rate k_on_=10^8^ M^-1^s^-1^ (42), the Ca^2+^ off rate from this site should not be faster than k_off_=K_d_k_on_=0.2 s^-1^ and the half-life of the bound state :23.5 s.

This raises an obvious question: what mechanism would allow the uniporter to bridge the ∼10^7^-fold difference in the Ca^2+^ dissociation rates? While mtCU might, arguably, represent an exceptional case of an ion channel, considering its function at micromolar external [Ca^2+^] and reliance on the membrane potential, the conundrum of reconciling the ion specificity with a nearly diffusion-limited transport rate is by no means unique to it. As originally proposed by Hodgkin and Keynes, for a potassium channel, there must be more than one ion binding site in the channel’s pore, and the ions must move through the pore in a single file (43). Consistent with this idea, numerous models of ion channels’ conductance have been considered over the years and tested through structural and computational studies (44–47).

Our results point to a mechanism that appears specific to MCU. The key difference, compared to other ion channels, is the extremely strong Ca^2+^ interaction with the selectivity filter of MCU. Our MD simulations suggest that the same ligands that trap Ca^2+^ in MCU’s selectivity filter also play an active role in transferring Ca^2+^ between two interaction sites in a process that we call the ligand-relay mechanism (Fig. 9). The four side chains of the E-ring (site 1) chelate Ca^2+^ tightly (N=7-8) yet have sufficient mobility to randomly detach and reach out for the incoming hydrated Ca^2+^ sequestered within the D-ring (site 2). This reduces the coordination number for the Ca^2+^ at site 1, lowering its binding energy and exposing it to the electrostatic field of the membrane potential, hence permitting its release and electrophoretic transport into the matrix. The Ca^2+^ conductance of MCU depends on the communication between the two sites, which requires structural flexibility within the DxxE motif. This mechanism differs from the knock-on mechanism of K^+^ transport through a potassium channel (48, 49). Here, the signature selectivity filter sequence TTVGYG is in an extended conformation, resulting in the main-chain carbonyl oxygens rigidly aligned within the channel’s pore. The K^+^ ions interact directly with each stationary site as they move in a single file down the selectivity filter (47). Different architecture is found in the pore of the voltage-gated sodium channels (50) and the structurally related calcium channels (51, 52). The pore comprises an outer funnel-like vestibule, a selectivity filter, a large central cavity, and the intracellular activation gate. Although a Glu residue, akin to that of the E-ring of MCU, occupies the central position in the selectivity filter, the opening is significantly wider than in MCU, permitting the conductance of Ba^2+^ or tetramethylammonium ions (44, 45). It does not appear capable of direct Ca^2+^ coordination, consistent with ∼1000x higher K_d_ for the Ca^2+^ block of Na^+^ conductance, and the transported ions remain partially hydrated at all times (53, 54).

**Figure 9.**
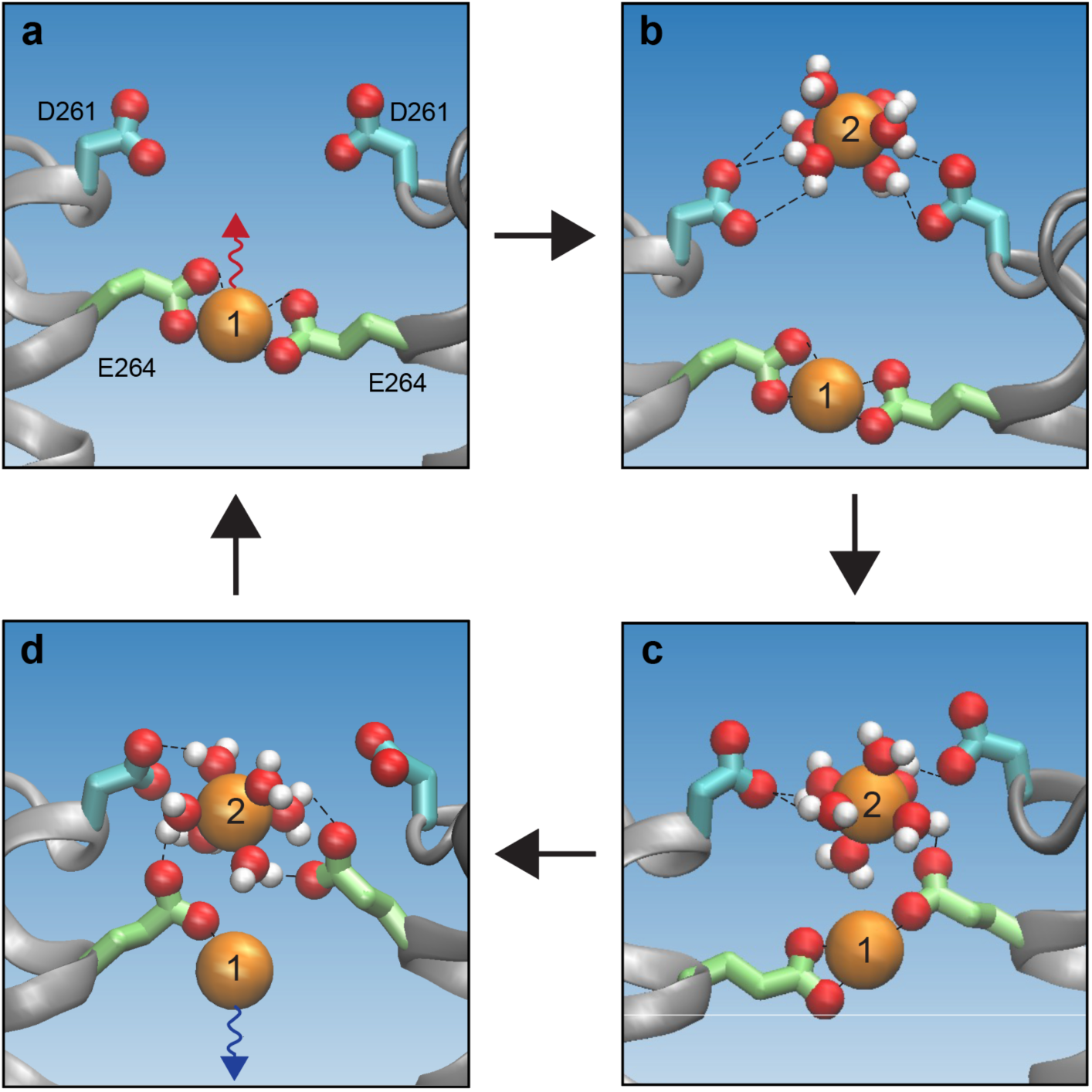
The proposed ligand-relay mechanism of the mitochondrial Ca^2+^ uptake via MCU. Shown is a cross-section of the channel in the DxxE region. For clarity of presentation, the atomic radii are not kept to scale. The blue-gradient background signifies the electrostatic field of the membrane potential (positively charged IMS on top). The wavy arrows symbolize the electrostatic forces acting upon the bound Ca^2+^. **A** – Bidentate coordination of the Ca^2+^ ion at site 1 by the four COO^-^ groups of the E-ring (only two are displayed). The complex (Ca^2+^(COO^-^)_4_) is pulled back towards the IMS site due to its negative charge. The channel is blocked. **B** – Sequestration of the incoming hydrated Ca^2+^ by the four Asp side chains of the D-ring (site 2). **C** – Transition of the ligands in the E-ring to a monodentate/bridging configuration. One carboxyl oxygen remains bound to the Ca^2+^ at site 1, while the other forms a hydrogen bond with the hydrated Ca^2+^ at site 2. In this “bridging” configuration, the negative charge of COO^-^ is distributed between site 1 and site 2. The electrostatic shielding effect of the E-ring for the Ca^2+^ bound at site 1 is reduced. **D** – One or more of the E-ring’s carboxyls become detached from the Ca^2+^ at site 1 and engage entirely with the Ca^2+^ at site 2. The electrostatic shielding for the Ca^2+^ ion at site 1 is lost, causing it to be propelled by the electrostatic field of the membrane potential toward the matrix site. The Ca^2+^ ion at site 2 loses its hydration shell and is carried by the attached Glu side chains of the E-ring into the site 1 position (as in panel A).

The mechanism we propose differs in key aspects from that published recently by Delgado and Long on the basis of their electrophysiological studies of WT and mutant *Tribolium castaneum* MCU-EMRE complexes reconstituted into planar lipid bilayers (14). In their experiments, the WT TcMCU-EMRE recapitulated the hallmark uniporter properties, such as the strong inward Ca^2+^ current rectification, distinctive ion selectivity, sub-nanomolar Ca^2+^ block of Na^+^ conductance, etc. All these properties were strongly affected by the E-to-D mutation in the E-locus of DxxE, whereas the D-to-A mutation in the D-locus was inconsequential, leading to the conclusion that the D-locus does not contribute to the ion conductance (14). This conclusion appears to be at odds with our model and with our and others’ observation that the D261A mutation in HsMCU reduces the mitochondrial Ca^2+^ uptake rate by a factor of >20 (cf. Table 1). These disparate conclusions might be reconciled if one considers the >1000x difference in [Ca^2+^] between the two types of experiments. At 40 mM [Ca^2+^], the conditions at which the electrophysiological recordings were performed, the diffusion-dependent direct interaction of one or more incoming Ca^2+^ ions with the E-locus may be sufficient to facilitate the release of the trapped Ca^2+^, effectively bypassing the D-locus and making the mutant’s conductance indistinguishable from that of the WT protein. However, under physiological conditions, the uniporter is exposed to only low micromolar [Ca^2+^]. At these conditions, diffusion-dependent random encounters between the incoming Ca^2+^ and that already bound in the selectivity filter are apparently insufficient, and transient sequestration of the incoming Ca^2+^ at the D-locus, as featured in our model, becomes essential.

Perhaps the most striking aspect of our proposed mechanism is its absolute dependence on the local structural dynamics of the DxxE motif and, possibly, of the entire TM-linker region. Importantly, this evolutionarily conserved feature is imparted by a residue not involved in Ca^2+^ binding. The contribution of local dynamics in enzyme catalysis has been studied extensively. It plays an important role in enzymatic turnover by enhancing the probability of sampling the transition state configuration (55, 56). The consequence of such an effect is that mutations at distant sites not directly involved in catalysis may have a profound effect on the enzyme kinetics. The switching off MCU activity by the P265A mutation in our studies is a vivid example of such an effect.

Our results suggest a potential mechanism of mtCU regulation that does not require physical blocking of the ion’s access to the pore or switching between distinct conformational states traditionally ascribed as the “on” and “off” states. The P265A mutation in HsMCU does not limit Ca^2+^ access to the pore, yet it is very effective in shutting off the channel’s conductance. The mutation appears to block the ligand-facilitated communication between the E-ring and the D-ring of DxxE, hence locking the selectivity filter with its bound Ca^2+^. Our results suggest that the uniporter’s function could be regulated kinetically through modulation of the TM-linker’s structural dynamics. It is conceivable that the MICU1/MICU2 complex might exert such an effect due to its contact with the D-ring or other segments of the TM-linker. It could also have a dual effect on the Ca^2+^ uptake rate, negative in the absence and possibly positive in the presence of Ca^2+^, hence permitting the switching of the uniporter on and off in step with cytosolic Ca^2+^ transients.

## Methods

### MCU sequence library and sequence conservation analysis

MCU homologs were identified by querying the human MCU (HsMCU) protein sequence (*Homo sapiens* Q8NE86) against the OrthoDB v. 10.1 (57), which linked to the InterPro database (IPR006769), release 80.1 (58). The retrieved sequences were aligned with MUSCLE (MUltiple Sequence Comparison by Log-Expectation) (32, 59) and manually curated by removing duplicates and sequences with large deletions or insertions in the region corresponding to the transmembrane domain of HsMCU. Also removed were excessively long sequences (>800aa). The final curated library contained 2495 MCU sequences. Using NCBI taxonomy identifiers and Entrez Programming Utilities, the taxonomical lineage was used to determine phylogenetic kingdom categories: *Metazoa, Viridiplantae*, and *Fungi*. All other species were placed into the “Other” category. Using an abbreviated version of the lineages with the 4 highest taxa levels, total species within the class taxon were counted, and the distribution was displayed using Krona (60). Sequence conservation was calculated with Consurf (33) (parameters: Bayesian calculation method, “Best model” for Evolutionary Substitution Mode). Adjusting the parameters to account for a maximum-likelihood tree reconstructed by IQ-Tree (61) did not drastically change our results. Residues that share interfaces with EMRE were identified by querying HsMCU (PDB: 6O58) on PDBePISA v. 1.52 (62).

### Secondary structure analysis

The main chain dihedral ϕ,ψ angles were computed from the respective structures using a PyMOL script. The following structures have been used: HsMCU (PDB: 6O58), fungal MCU (PDB: 6C5W, 6D7W), and human uniporter holocomplexes (PDB: 6K7X, 6WDN, 6WDO, 6XJV, 6XJX). The Ramachandran plot and the ϕ^2^,ψ^2^plot for each structure were generated for the TM-linker region and the flanking transmembrane helices TM1 and TM2, as defined by the structural and sequence alignments. For the corresponding sequence representation of the ϕ^2^,ψ^2^ plots, their values were averaged for all eight MCU chains in each structure, and the baseline values corresponding to the membrane-embedded sections of TM1 and TM2 were subtracted.

### Structure preparation with hexaamminecobalt(III)

Using previously derived hexaamminecobalt(III) parameters (NCO) (63), the CHARMM parameter and topology files were generated using ForConX (64). To generate WT-HsMCU+NCO, a CeMCU-EMRE model (PDB: 6DNF, chains A to H), which contains 2 Ca^2+^ ions bound within the channel’s pore, was aligned with HsMCU (PDB: 6O58). The Ca^2+^ ion located within the D-ring was replaced with NCO.

### Molecular dynamics simulations

The Membrane Builder in CHARMM-GUI server (65–67) was used to prepare the cryo-EM structure of HsMCU (PDB: 6O58, residues 216-310) for MD simulations. Hydrogen atoms were added to the protein, and the protein was inserted into a preassembled lipid bilayer with 1-palmitoyl-2-oleoylphosphatidylcholine (POPC) lipids. A 22.5 Å water layer was on both sides of the lipid bilayer. The cubic box of ∼100 × ∼100 × ∼110 Å contained 55 potassium and 75 chloride ions to neutralize the simulation box at neutral pH and for an ionic strength of 150 mM. The final system contained one Ca^2+^ ion coordinated by Glu264 sidechains of the four MCU chains, with or without NCO, and ∼20600 TIP3 water molecules, for a total of ∼100500 atoms.

All MD simulations were performed in NAMD (68) using the CHARMM force field (v. CHARMM36m) parameters (69), and visualized in VMD. The system was equilibrated at constant temperature (310 K) and constant pressure (1.0135 atm) by employing Langevin dynamics and Langevin piston (70, 71), with the protein-backbone atoms, Ca^2+^ ion, and NCO ligand, if applicable, were harmonically constrained to their initial positions and then slowly released using a decreasing force constant equal to 10; 5; 2.5; 1; 0.5; and 0.1 kcal/mol/Å^2^. After the backbone was completely released, we ran an unconstrained simulation for 20 ns, with the position coordinates saved to the trajectory file every 0.1 ns for 200 frames. The RMSD Trajectory Tool plugin in VMD was used to assess the structural stability during the MD simulations.

### Analysis of MD simulation data

The secondary structure distribution was analyzed with the Timeline plugin of VMD, using the default definitions of the components encoded in VMD according to Structural Identification (STRIDE) criteria for T: turn, H: α-helix, G: 3_10_ helix, C: coil, I: π-helix, E: extended conformation, B: isolated bridge. The distribution of secondary structure types across residues 252-265 was calculated and plotted using the Pandas package for Python3 in Jupyter Notebooks. Backbone H-bond interactions were determined for a given residue *i* by computing the distance and angle between residue *i*→*i*+3 (3_10_ helix); *i*→*i*+4 (α-helix); and *i*→*i*+5 (π-helix) using an in-house generated tcl script. We applied a distance cutoff of 3.6 Å and angle cutoff 32°, where distance is defined as donor atom (D) and acceptor atom (A) distance, and angle is defined as the A-D-H angle. Violin plots for C-alpha distances were constructed by measuring the distances between the C_α_ atoms of a given residue *i* for 6 residues (Trp260, Asp261, Ile262, Met263, Glu264, Pro265/Ala265), and its corresponding *i+3* and *i+4* residue was calculated for WT-HsMCU and P265A-HsMCU trajectories. Violin plots were generated with python 3.

### HEK293T cells culture and double knock-out cell line production

HEK293T cells were cultured in Dulbecco’s Modified Eagle Medium (DMEM) (Life Technologies, 11995) with 10% FBS, penicillin/streptomycin S, and 1X GlutaMax (Gibco). The HEK293T MCU knock-out cell line was generated using the TALE nuclease (TALEN) (6). The double knock-out cell line was generated by CRISPER/Cas9 system to knock-out MCUb in MCU knock-out cell line. The HsMCUb sgRNAs were cloned into pSpCas9(BB)-2A-GFP (Addgene #48138) plasmid (72). The plasmids had been transfected to cells by Lipofectamine 2000. After three days of transfection, the GFP-positive cells were sorted by FACS into a 96-well plate for clonal isolation. The MCU/MCUb KO cell clones were verified by sequence analysis, HsMCUb sgRNA: GTCACACCATTATAGTACCG.

### Cloning, mutagenesis, virus production, and cell transduction

The cDNA of MCU was cloned into pLYS5 (Addgene, #50054) with a FLAG tag at the C-terminus (6). The point mutation plasmids were generated using QuikChange II kit (Agilent, cat no. 200521) with corresponding primers. All the clones have been confirmed by sequencing using CMV-forward primer, and custom reversed primers. Lentivirus for each mutant was generated by mixing 100 ng of VSV-G, 900 ng of psPax2, and 1μg of the viral plasmid into 200 μl of DMEM supplied with 6 μl of X-tremeGENE™ HP and incubating at room temperature for 20 min. Then the virus mixture was added to 1 million HEK293T cells in a 6-cm dish with 5 ml of culture medium. After two days of transfection, the supernatant of the culture medium, which contains the virus, was filtered with 0.45µm sterile PES filter and stored at -80°C. The 250K HEK293T cells were transduced with 200 μl of virus-containing solution on a 6-well dish with 8 µg/ml of polybrene in 2 ml of culture medium. After two days of transduction, the virus medium was removed. The selection was made by adding fresh medium with 2µg/ml of puromycin dihydrochloride and/or 100µg/ml of hygromycin B.

### Calcium uptake measurements

The calcium uptake experiments were performed in 96 well plates with 1 million cells for each sample. The cells were permeabilized in KCl buffer (125 mM KCl, 2mM K_2_HPO_4_, 1mM MgCl_2_, 20 mM HEPES at pH 7.2, 0.005% digitonin, 5 mM glutamate, 5 mM malate, and 5 µM EGTA) with 1 µM of Calcium Green™-5N (Invitrogen). 30 µM of CaCl_2_ was injected into wells at a specific time. The Ru360, when present, was added to samples at 1µM. The fluorescence signal was recorded with a PerkinElmer Envision plate reader with excitation at 485 nm and emission at 535 nm. Calcium uptake rate constants were calculated using single exponential decay to fit the curve from the highest fluorescence signal to the baseline (GraphPad). The calcium uptake inhibition experiments with Hexaamminecobalt(III) (NCO) were performed in KCl buffer without EGTA. Appropriate concentrations of NCO were added to each well before the addition of CaCl_2_.

### Protein sample preparation, gel electrophoresis, and Western blots

The cells were harvested in lysis buffer (50 mM HEPES, 100 mM NaCl and 1% of TritonX-100, pH=7.4) with protease inhibitors on ice. The lysis samples were centrifuged at 17,000xg for 10 min. The supernatant was taken for Bradford assay to determine the protein concentration. The protein samples were loaded and run in 4-20% SDS-PAGE. Then the gel was transferred on the PVDF membrane with a Trans-Blot Turbo system (Bio-Rad) with mix molecular weight protocol. After transfer, the PVDF membrane was washed in 1X TBST and incubated with 5% of milk in TBST for 30min. The list of antibodies and dilution has been used in this study: MCU (Sigma-Aldrich, Cat. no. HPA016480, 1:1,000), ATP5A (Abcam Biochemicals, Cat. no. ab14748, 1:5,000). The antibodies were diluted in 5% milk TBST and membrane incubated in a cold room overnight. Then, the membrane was washed in TBST for 5 min three times and blotted with HRP-linked secondary antibodies at room temperature for 1 hour. The membranes were developed in ECL substrate (PerkinElmer, Cat. no. ORT2755, ORT2655) and imaged by ChemiDoc MP system (Bio-Rad).

Primers for mutagenesis:

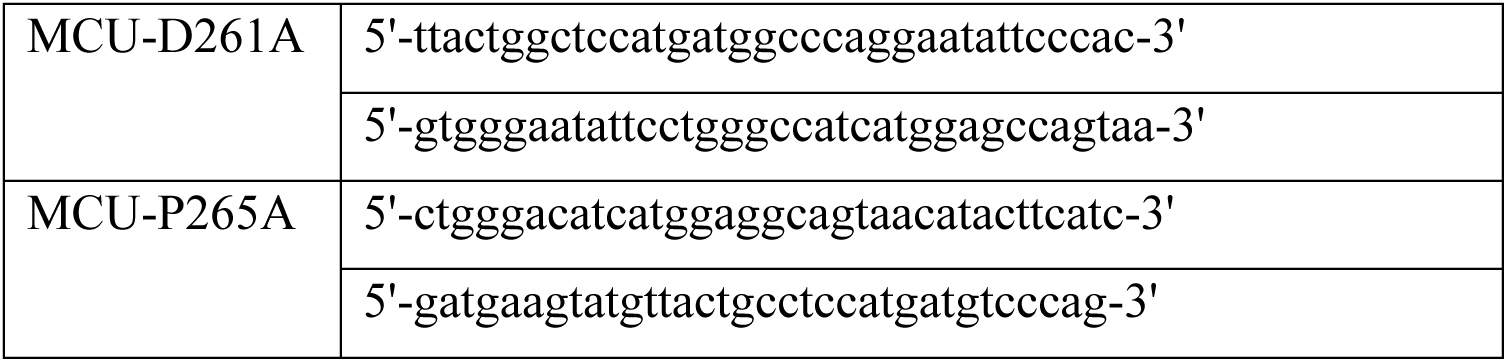

## ACKNOWLEDGMENTS

We thank Vamsi K. Mootha for his support, advice, and critical comments on the manuscript. We thank Stefan Boresch and Christian Schröder for their advice on generating the CHARMM-compatible NCO parameters using ForConX. We thank Ruma Banerjee and Rohit Sharma for their insightful comments on the manuscript. This work was supported by National Institutes of Health Grants R01AR071942 (to Z.G.) and K08HL157620 (to P.S.W.).

## Supplementary Information

Figures S1-S5, Table S1

**Figure S1.**
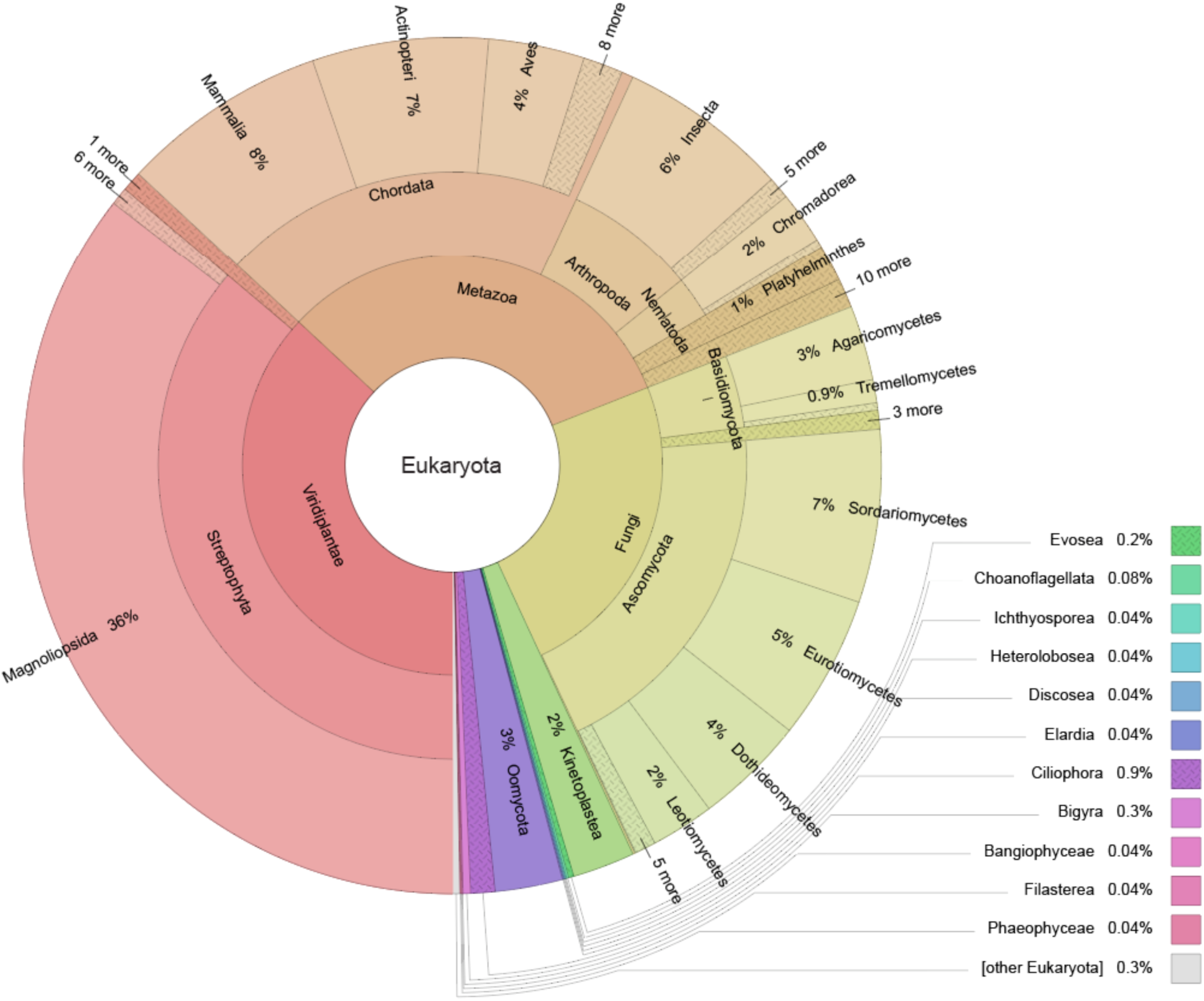
Distribution of species in the MCU sequence library used for the sequence conservation analysis. An abbreviated form of the lineages is used, displaying the domain, kingdom, phylum, and class taxa levels. This graph was generated using Krona: https://bio.tools/krona

**Figure S2.**
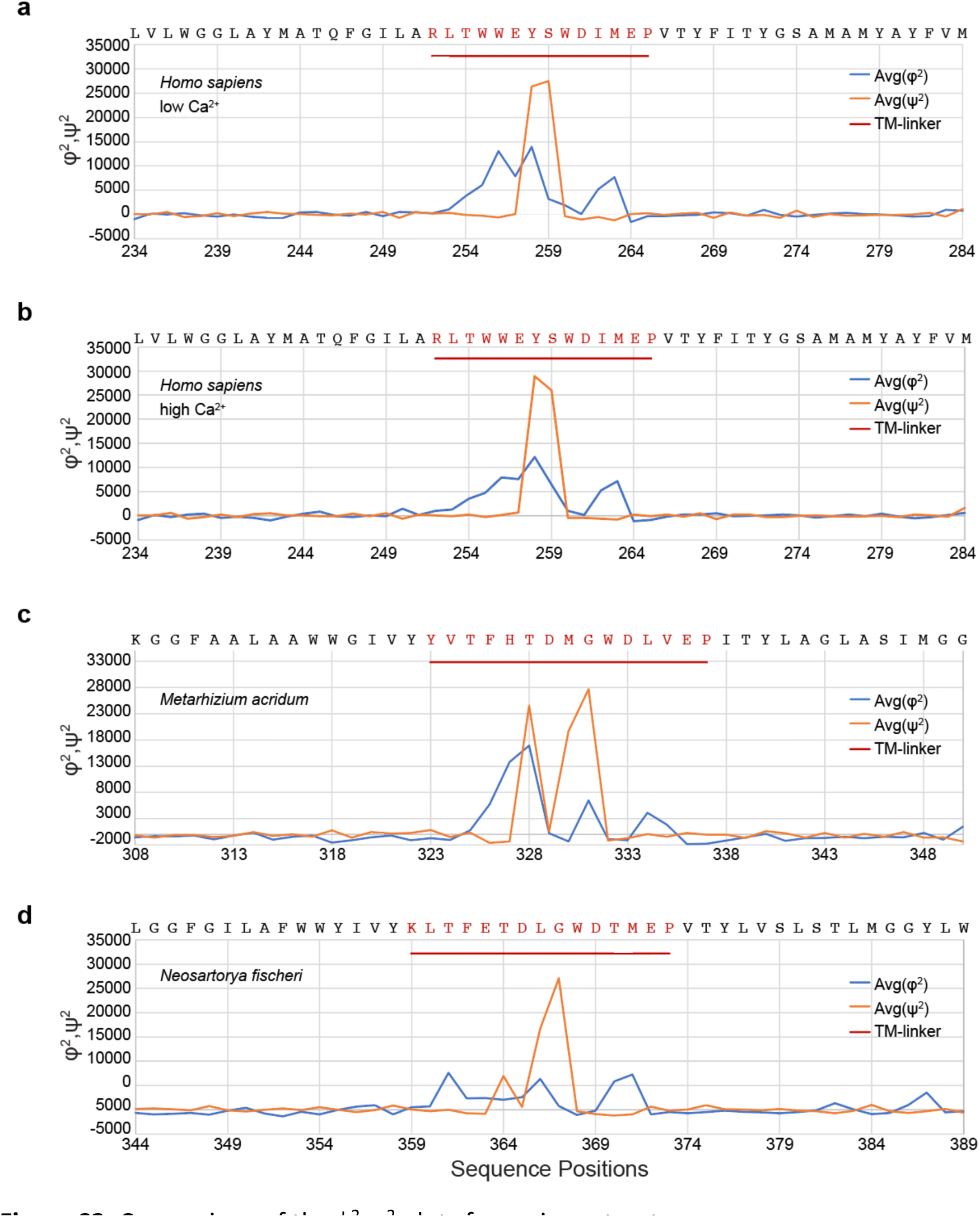
Comparison of the ϕ^2^,ψ^2^ plots for various structures. **A** - Human uniporter holocomplex in low Ca^2+^. PDB:6XJX. **B** - Human uniporter holocomplex in high Ca^2+^, PDB:6XJV. **C** - *Metarhizium acridum* MCU, PDB:6C5W. **D** - *Neosartorya fischeri* MCU, PDB:6D7W. The TM-linker highlighted in red.

**Figure S3.**
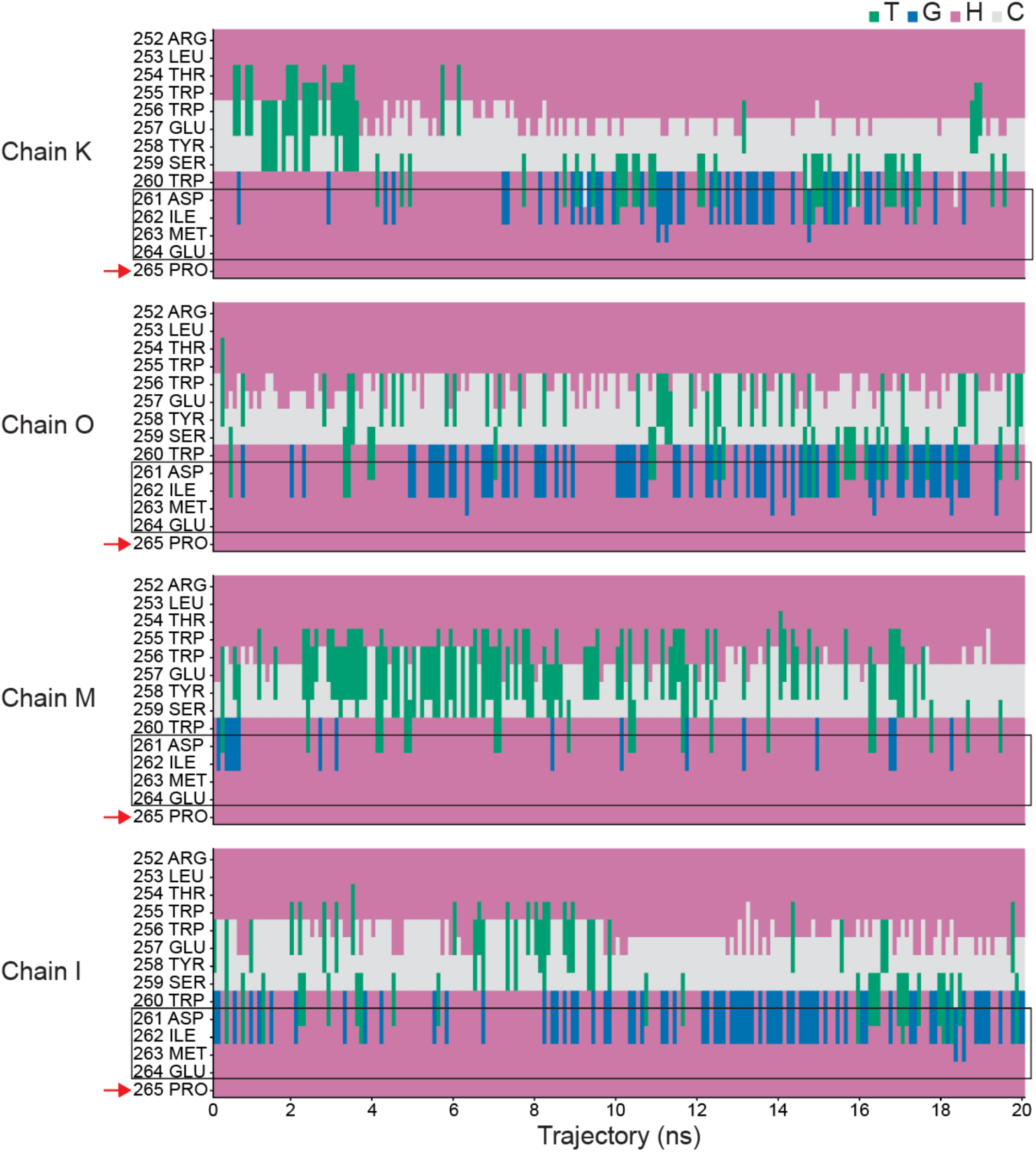
Secondary structure fluctuations in the TM-linker of WT-HsMCU. All four chains of the tetrameric MCU complex are shown (PDB:6O58). The DxxE motif is outlined with a rectangle. Color coding for the secondary structure types: Green – turn (T), Blue-3_10_-helix (G), Pink – α-helix (H), Grey – extended/coil (C)

**Figure S4.**
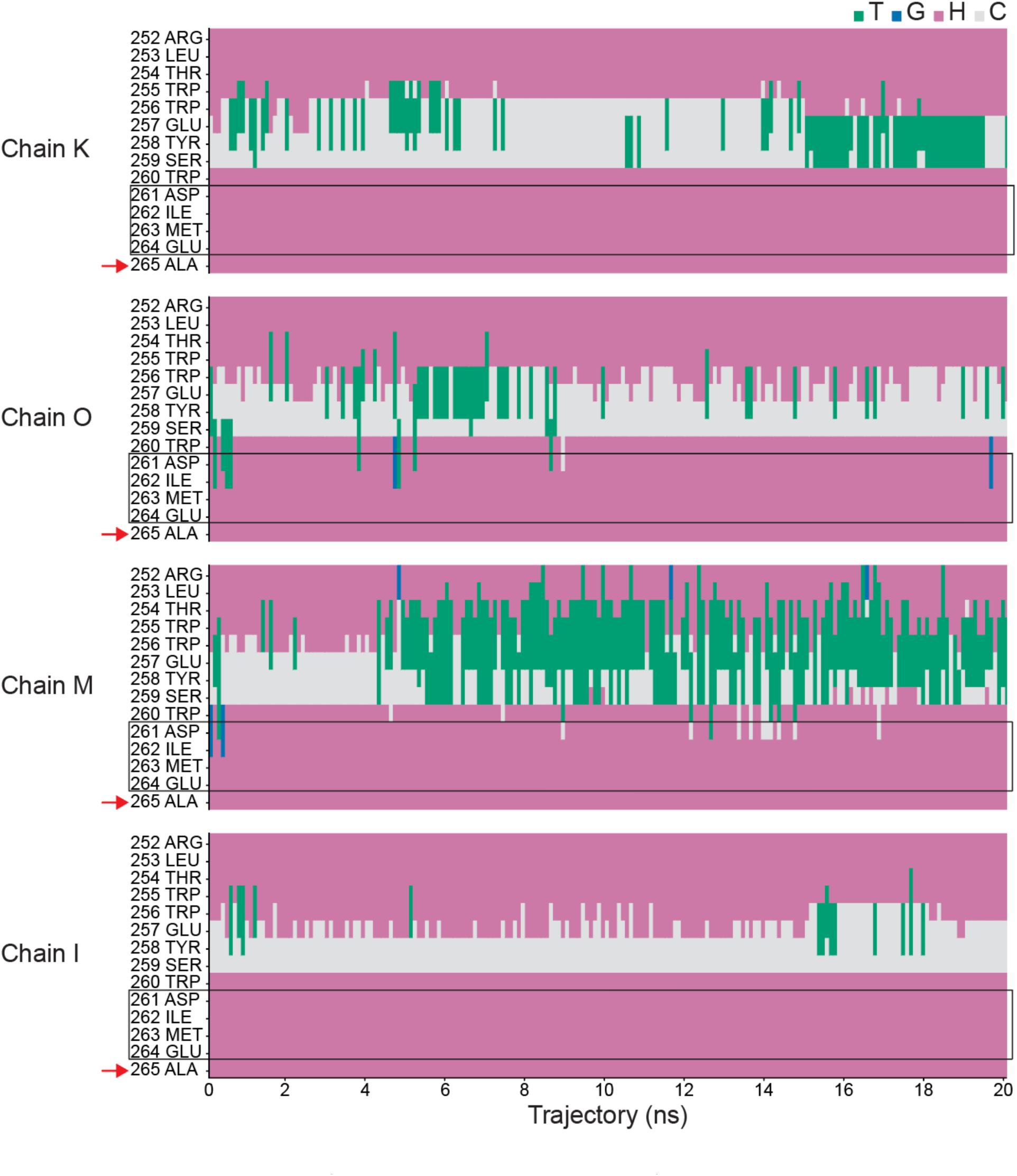
Secondary structure fluctuations in the TM-linker of the P265A-HsMCU mutant. The secondary structure designation is the same as in Suppl. Fig. 3. The red arrow indicates the position of the mutation. Note the absence of secondary structure fluctuations in the DxxE region (outlined with a rectangle).

**Figure S5.**
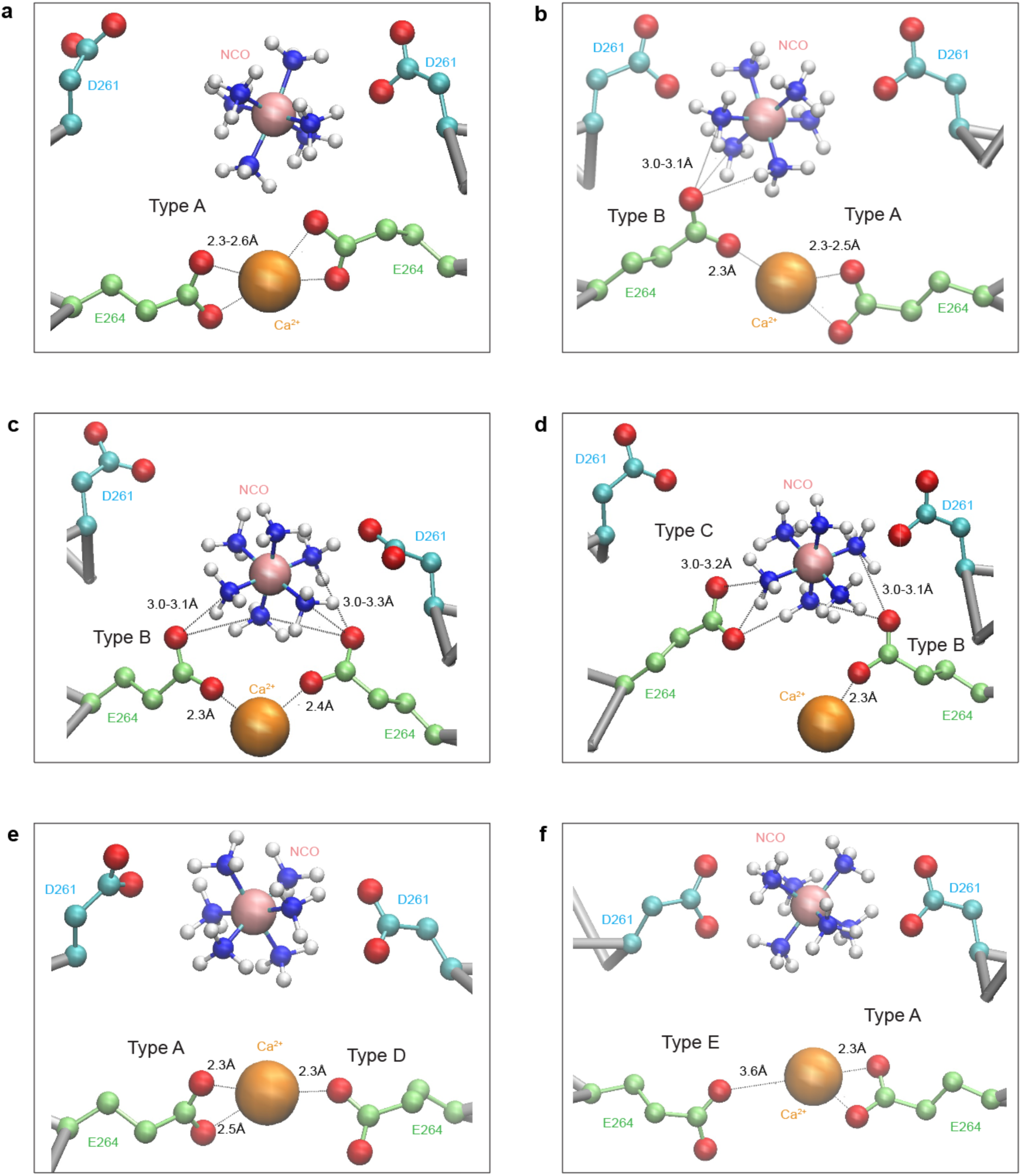
Examples of various types of interactions in the E-locus of MCU. Shown are representative snapshots of the channel cross-section in the DxxE region taken from a 20 ns MD simulation of WT-HsMCU with bound Ca^2+^ and NCO. See Fig. 8 in the main text for a definition of the five most frequent types of interactions labeled here as Type A – Type E.

**Table S1.** The distorted geometry of the TM2 helix. Mean values of C_α_-C_α_ distances and their standard deviations are shown (n=800, four polypeptide chains of MCU tetramer averaged over 20 ns of molecular dynamics simulation).

| Residue | WT-HsMCU |  | P265A-HsMCU |  |
| --- | --- | --- | --- | --- |
|  | C<sub>α</sub>-C<sub>α</sub> i,i+3 (Å) | C<sub>α</sub>-C<sub>α</sub> i,i+4 (Å) | C<sub>α</sub>-C<sub>α</sub> i,i+3 (Å) | C<sub>α</sub>-C<sub>α</sub> i,i+4 (Å) |
| Trp260 | 4.80 ± 0.23 | 5.85 ± 0.36 | 4.89 ± 0.25 | 5.82 ± 0.28 |
| Asp261 | 5.71 ± 0.34 | 7.81 ± 0.36 | 5.57 ± 0.37 | 6.70 ± 0.54 |
| Ile262 | 6.56 ± 0.40 | 7.06 ± 0.42 | 5.35 ± 0.39 | 6.37 ± 0.37 |
| Met263 | 5.05 ± 0.26 | 6.02 ± 0.29 | 5.05 ± 0.26 | 6.08 ± 0.22 |
| Glu264 | 5.31 ± 0.24 | 6.30 ± 0.27 | 5.05 ± 0.25 | 6.18 ± 0.28 |
| Pro/Ala265 | 5.00 ± 0.21 | 6.12 ± 0.21 | 5.13 ± 0.21 | 6.32 ± 0.25 |
| Val266 | 5.12 ± 0.21 | 6.26 ± 0.22 | 5.12 ± 0.26 | 6.25 ± 0.21 |
| Thr267 | 5.16 ± 0.23 | 6.16 ± 0.24 | 5.13 ± 0.23 | 6.10 ± 0.20 |

